# Investigation of the usefulness of liver-specific deconvolution method toward legacy data utilization

**DOI:** 10.1101/2023.04.19.537436

**Authors:** Iori Azuma, Tadahaya Mizuno, Katsuhisa Morita, Yutaka Suzuki, Hiroyuki Kusuhara

## Abstract

**Background:** Immune responses in the liver are related to the development and progression of liver failure, and precise prediction of their behavior is important. Deconvolution is a methodology for estimating the immune cell proportions from the transcriptome, and it is mainly applied to blood-derived samples and tumor tissues. However, the influence of tissue-specific modeling on the estimation results has rarely been investigated. In this study, we constructed a system to evaluate the performance of the deconvolution method on liver transcriptome data.

**Results:** We prepared seven mouse liver injury models using small-molecule compounds with known hepatotoxicity and established a dataset with corresponding liver bulk RNA-Seq and immune cell proportions, covering various immune responses. RNA-Seq expression for nine leukocyte subsets and four liver-associated cell types were obtained from the Gene Expression Omnibus (GEO) to provide a reference covering liver component cells. Here, we found that the combination of reference cell sets affects the estimation results of reference-based deconvolution methods. We established a liver tissue-specific deconvolution by optimizing the reference cell set for each cell to be estimated. We applied this model to independent datasets and showed that liver-specific modeling focusing on reference cell sets is highly extrapolatable.

**Conclusions:** We provide an approach of liver-specific modeling when applying reference-based deconvolution to bulk RNA-Seq data and show its importance. It is expected to enable sophisticated estimation from rich tissue data accumulated in public databases and to obtain information on aggregated immune cell trafficking.

## Background

- The deconvolution method extracts immune cell information, such as the proportion of immune cells in the sample from the bulk transcriptome data. The bulk transcriptome has a long history, starting with microarrays around 2000, and much data have been accumulated in public databases and are easily available [1]. Therefore, combining the deconvolution method with these legacy data is expected to enable us to obtain aggregated knowledge on the behavior of various immune cells, so-called immune cell trafficking, under various conditions. Acquiring such comprehensive insights is challenging with flow cytometry data, which lacks a well-structured database. While repositories for scRNA-seq data have emerged, their quantity remains limited in comparison to bulk RNA-Seq. Additionally, the expenses associated with utilizing public databases, including reprocessing costs, remain notable [2].

Typical deconvolution methods are reference-based methods that use the unique gene expression levels of immune cells as prior information [3–5]. When these methods are applied to tissue data accumulated as legacy data, how accurate is the estimation? Whereas evaluation datasets exist for blood, and their performance has been evaluated [3, 6–8], is there any tissue dependence when applying these methods to tissues with other parenchymal cells? Such a possibility has been highlighted by Chen and Wu. [9]. However, because no evaluation dataset exists, there was no clear answer.

In the present study, we analyzed mouse liver tissues to verify the accuracy of the reference-based deconvolution method in tissues. Immune cell trafficking, especially that of neutrophils, is important in the development and progression of liver failure, and studies using various liver injury models have reported the role of immune cells in the migration of neutrophils [10–14]. However, the studies have been limited to individual models of specific disorders, and there is no aggregated knowledge of the commonalities and differences in the contribution rate and the order of elicitation of each cell type.

Therefore, we evaluated the accuracy of the deconvolution method by establishing an evaluation dataset covering various immune cell behaviors using compound-induced liver injury models, in which there is relatively little confounding among the models.

The present study has three contributions to the understanding of immune responses in tissues using deconvolution, as follows.

(1) Using several liver injury models induced by each of seven small-molecule compounds with known hepatotoxicity, we constructed a dataset to evaluate the deconvolution method covering various immune responses.
(2) In the reference-based deconvolution method, we showed that selecting cell types that constitute the reference is important.
(3) By using a liver-specific optimized model, we found behaviors of immune cells that could not be found by conventional methods.

## Results

### Establishment of the evaluation dataset using drug-induced liver injury models

No evaluation dataset corresponds to the liver sample transcriptome and immune cell type proportions. To construct a liver-specific deconvolution model, we first prepared an evaluation dataset that reflects the immune response in liver tissue. In general, the evaluation dataset should be as diverse as possible regarding input–output relationships and free of confounding factors other than inputs and outputs. We prepared a diverse and low confounding dataset using perturbation by various small molecules. Liver injury models with small molecular compounds are expected to contribute to the establishment of a highly diverse dataset, with little confounding by procedures such as surgery. Based on a literature survey, we selected seven compounds, alpha-naphthyl isothiocyanate (ANIT), acetaminophen (APAP), CCl4 (CCl4), concanavalin A (ConA), galactosamine (GAL), 4,4′-methylene dianiline (MDA), and thioacetamide (TAA), that are frequently used to induce liver injury [15–21].

The administration of each compound-induced liver injury, and tissues were collected 24 h after administration. Flow cytometry, RNA-Seq analysis, and blood biochemistry tests were performed on the obtained tissues, and an evaluation dataset was established in which these measurements corresponded to each sample (Fig. 1a).

**Fig. 1.**
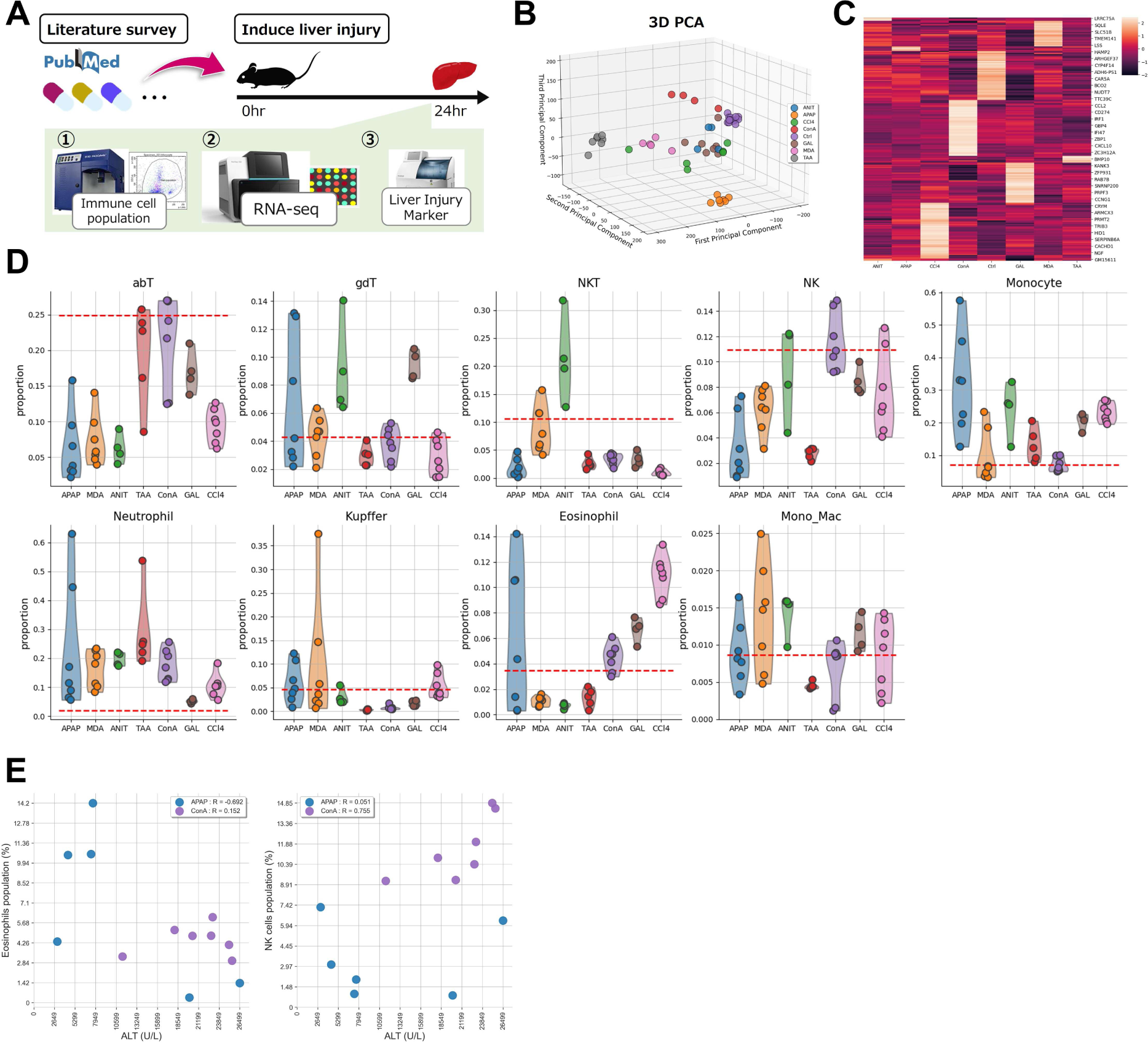
Measurement of transcriptome and immune cell proportions using drug-induced liver injury mouse models. **A** We selected 7 compounds with known hepatotoxicity and administered them to mice to induce liver injury. After 24 h of administration, liver was harvested for flow cytometry and RNA-Seq analysis, and blood was collected to measure the liver injury markers. **B** PCA plotting of the obtained RNA-Seq data shows clusters formed by each compound administration. **C** Heatmap showing median gene expression level (*z* score) of up to 50 differentially expressed genes in each administration group. **D** Violin plots showing the proportion of the nine immune cell subtypes to the CD45^+^ cell population in each liver injury sample was measured by flow cytometry. The red dashed line indicates the control samples without perturbation. **E** Scatterplot shows the Pearson correlation between alanine aminotransferase (ALT) values and immune response. The population of eosinophils after APAP administration correlated negatively with the ALT values, and NK cells after ConA administration correlated positively with the ALT values.

### RNA-Seq analysis of the evaluation dataset

Liver transcriptome data were obtained by RNA-Seq. We performed PCA on the processed data and showed that each treatment group formed a cluster (Fig. 1b). The average expression levels of the genes in each treatment group were calculated and visualized in a heatmap, which detected differentially expressed genes (DEGs) among the treatment groups (Fig. 1c). These results indicate that each of the seven compounds used to induce liver injury had specific effects on the tissue and that gene expression profiles were separated among the treatment groups. Thus, our established evaluation dataset using drug-induced liver injury models is expected to reflect the diverse immune responses in the liver (Fig. S1).

### Flow cytometry analysis on the evaluation dataset

We analyzed the flow cytometry data with FlowJo software. In the present study, we obtained ground truth data on the proportion of nine subsets representative of the liver, αβT cells (CD45^+^/CD3^+^/gdTCR^−^), γδT cells (CD45^+^/CD3^+^/gdTCR^+^), natural killer T (NKT) cells, natural killer (NK) cells (CD45^+^/CD3^+^/NK1.1^+^), monocytes (CD45^+^/CD11b^+^/Siglec-F^−^/Ly6G^−^/Tim4^−^/Ly6C^+^), neutrophils (CD45^+^/CD11b^+^/Siglec-F^−^/Ly6G^+^/Tim4^−^), Kupffer cells (CD45^+^/CD11b^+^/Siglec-F^−^/Ly6G^+^/Tim4^+^/Ly6C^−^/F4/80^+^), eosinophils (CD45^+^/CD11b^+^/Siglec-F^+^), and monocyte-derived macrophages (CD45^+^/CD11b^+^/Siglec-F^−^/Ly6G^−^/Tim4^−^/Ly6C^−^), in compound-induced liver injury samples (Fig. S2, Supplementary Note). Each immune cell can be evaluated for increase or decrease compared with untreated control samples. We observed some characteristic immune cell trafficking for each compound administration (Fig. 1d). For instance, ConA administration led to an increase in monocytes, while TAA administration resulted in a decrease in NK cells. Furthermore, in response to APAP administration, there was significant variability in the immune response across individuals, with certain samples exhibiting elevated levels of neutrophils and monocytes. These results were consistent with existing findings [22–25]. To our knowledge, this is the first report of an increase in eosinophils after GAL administration, and the responses of various immune cells after MDA administration were captured using flow cytometry.

Notably, in the present analysis, each sample was tied to the value of liver injury markers so that we could associate the degree of damage with the behavior of immune cells (Fig. S3). There were large individual differences in liver injury in the APAP and ConA treatment groups, which could be stratified by the accumulation of immune cells. Specifically, the trafficking of eosinophils after APAP administration varied widely, but there was a tendency for eosinophils not to accumulate in the severely injured samples with high alanine aminotransferase (ALT) and aspartate aminotransferase (AST) levels. Furthermore, ALT and AST values at ConA administration correlated with the proportion of NK cells and neutrophils (Figs 1e and S4). In the present study, the established evaluation dataset was used to evaluate and optimize the deconvolution method, but we would also like to emphasize that it provides novel insights useful for understanding the mechanisms underlying the induction of liver injury and immune cell trafficking by each compound administered.

### Evaluation of the impact of the reference cell sets on estimation performance

The immune cell proportions were estimated by deconvolution using Elastic Net on the established evaluation dataset. In the present study, we obtained cell-specific gene expression profiles for up to 13 types of cells derived from immune cells and liver cells and made them available as references (Table 1). The correlation and multicollinearity of the gene expression profiles of these 13 cell types are shown in Figure S5.

**Table. 1.** Summary of sample information used to construct a reference consisting of up to 13 cell types.

| Cell / Dataset | PubMed ID | Platforms | Selected Samples | Original Samples | Criteria |
| --- | --- | --- | --- | --- | --- |
| B cells | - | - | 11 | - | - |
| GSE84878 | 28538178 | Illumina HiSeq 2000 | 7 | 7 | WT |
| GSE61425 | 25288398 | Illumina HiSeq 2000 | 4 | 4 | WT |
| CD4 + T cells | - | - | 13 | - | - |
| GSE119852 | 31253760 | Illumina HiSeq 4000 | 4 | 4 | Blood control |
| GSE152757 | 32839313 | Illumina NovaSeq 6000 | 9 | 9 |  |
| CD8 + T cells | - | - | 14 | - | - |
| GSE151936 | 32814898 | Illumina NextSeq 500 | 12 | 15 | Spleen |
| GSE70813 | 27102484 | Illumina HiSeq 2500 | 2 | 4 | Spleen |
| NK cells | - | - | 8 | - | - |
| GSE114827 | 30127438 | Illumina NextSeq 500 | 3 | 3 | Liver WT |
| GSE103901 | 29056343 | Illumina HiSeq 2500 | 5 | 5 | Spleen |
| Neutrophils | - | - | 10 | - | - |
| GSE142432 | 33098771 | Illumina HiSeq 2500 | 6 | 6 | Blood, Spleen |
| GSE116177 | 30395284 | Illumina HiSeq 2500 | 4 | 4 | Peripheal Blood |
| Eosinophils | - | - | 7 | - | - |
| GSE55385 | 25765318 | Illumina HiSeq 2000 | 4 | 4 |  |
| GSE145985 | - | Illumina NovaSeq 6000 | 3 | 3 |  |
| Basophils | - | - | 3 | - | - |
| GSE132122 | 33547048 | Illumina NextSeq 500 | 3 | 3 |  |
| Monocytes | - | - | 6 | - | - |
| GSE130257 | 31444235 | Illumina NextSeq 500 | 2 | 2 | Blood |
| GSE116177 | 30395284 | Illumina HiSeq 2500 | 4 | 4 | Peripheal Blood, Bone Marrow |
| Kupffer cells | - | - | 12 | - | - |
| GSE152211 | 34469774 | Illumina NovaSeq 6000 | 6 | 6 |  |
| GSE138778 | 32562600 | Illumina NovaSeq 6000 | 6 | 6 | Chow diet |
| Hepatocytes | - | - | 10 | - | - |
| GSE104415 | 29618815 | Illumina NextSeq 500 | 6 | 6 |  |
| GSE132368 | 32001510 | Illumina NextSeq 500 | 4 | 4 | DMSO 0hr |
| Cholangiocytes | - | - | 3 | - | - |
| GSE156894 | - | NextSeq 550 | 3 | 3 |  |
| Stellate cells | - | - | 6 | - | - |
| GSE135789 | 31561945 | Illumina NextSeq 500 | 3 | 4 |  |
| GSE120281 | 34277612 | Illumina HiSeq 3000 | 3 | 3 |  |
| LSEC | - | - | 10 | - | - |
| GSE120281 | 34277612 | Illumina HiSeq 3000 | 3 | 3 |  |
| GSE120282 | 34277612 | Illumina HiSeq 3000 | 3 | 3 | Control |
| GSE135789 | 31561945 | Illumina NextSeq 500 | 4 | 4 | WT |

We prepared three representative reference cell sets. The first is a set of six cell types called LM6 (neutrophils, monocytes, B, CD8, CD4, and NK), which are frequently used in estimating the proportion of immune cells in blood using the deconvolution method [25, 26]; the second is LM9, to which we added immune cells that are not frequently used but are important in the liver (e.g., eosinophils, basophils, and Kupffer cells); and the third is LM13, to which we added four cell types related to the liver. Using these references, we evaluated the estimation performance on the evaluation dataset (Fig. 2a–c).

**Fig. 2.**
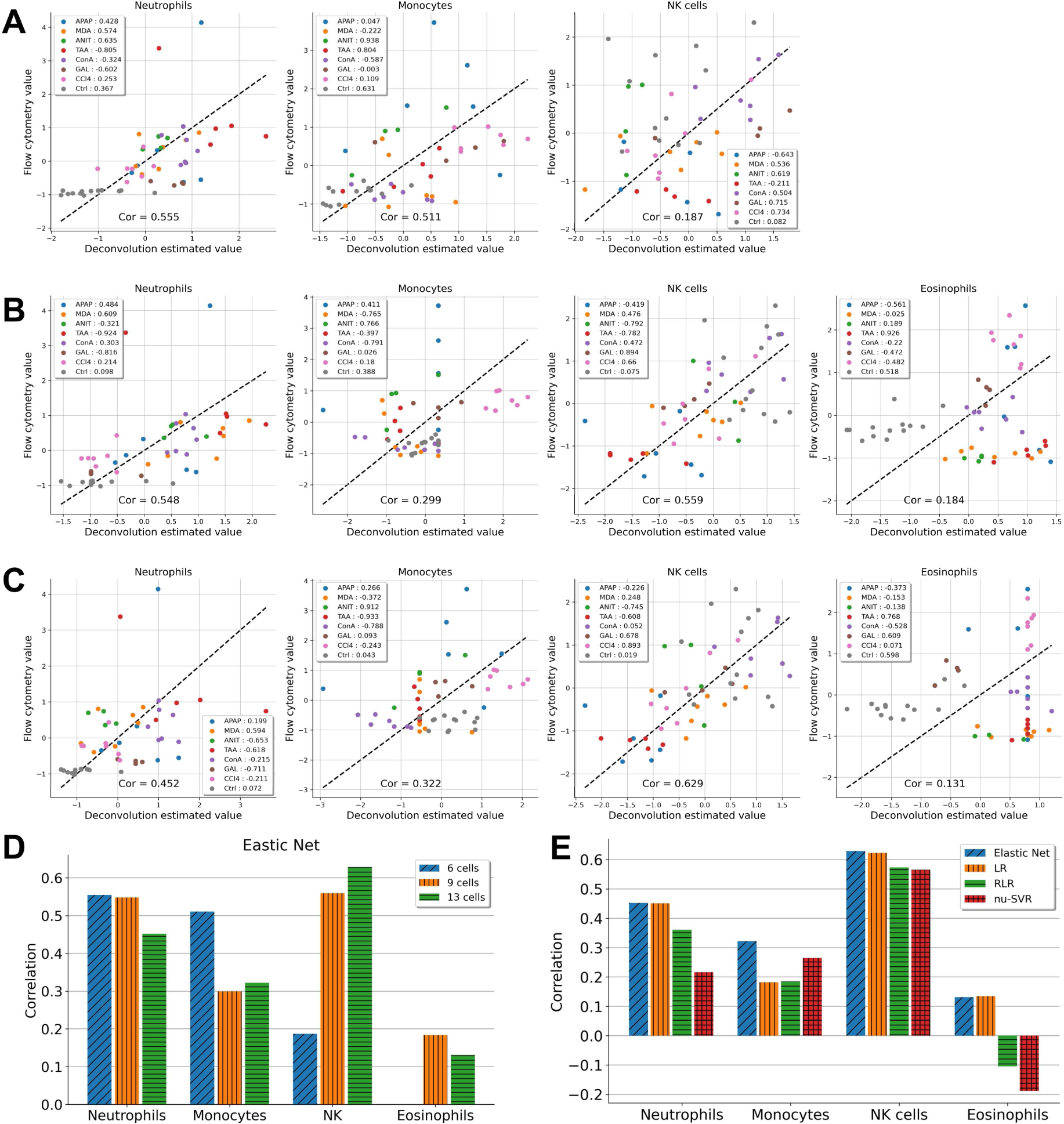
Conducting a reference-based deconvolution method on the dataset for evaluation. The scatterplot shows the Pearson correlation between flow cytometry values and deconvolution estimates using Elastic Net when **A** LM6, **B** LM9, and **C** LM13 are used as reference. **D** Bar plots showing the summary of the impact of the reference cell set on estimation performance. Eosinophils are not included in LM6 and cannot be estimated when 6 cell types are used as reference. **E** Bar plots showing the estimation performance of the representative methods when using all 13 cell types.

To estimate neutrophils, LM6 showed superior performance, and LM13 showed poor performance. In contrast, to estimate NK cells, LM6 performed poorly, but LM13 performed well. The reference cell type dependence was also confirmed for other representative algorithms other than Elastic Net (Fig. S6). Additionally, these findings were replicated in the analysis utilizing scRNA-Seq data obtained from a single experiment as a reference (Figs S7a–c). These findings indicate that the estimation accuracy of each cell depends on the combination of cell sets placed in the reference and that there is no reference with consistently good estimation performance (Fig. 2d and Fig. S7c). Conversely, assessments with artificial pseudo-bulk datasets simulating the liver consistently demonstrated elevated estimation performance for LM6, LM9, and LM13 (Fig. S8). These outcomes imply that the influence of the reference cell set is a manifestation of the nonlinear attributes inherent in real-world data. It is noteworthy that analyses employing pseudo-bulk datasets, constructed under rigorous linearity-preserving constraints, might lead to an overestimation of deconvolution methods. Consequently, we selected Elastic Net, which exhibited superior performance with all 13 cell types, and fine-tuned this approach for subsequent analyses (Fig. 2e).

### Optimization of the combination of reference cell types

In general, deconvolution is the methodology used to estimate the composition of cells. However, when describing individual tissue conditions, such as elucidating disease mechanisms or stratification, it is important to focus on a specific cell type and estimate how the cell fluctuates compared with the control group in terms of trafficking. In the case of trafficking, building a model for each cell is possible because it is specified for a control group. Therefore, we optimized the model for each target cell and evaluated the characteristics of the reference cell sets.

When estimating the trafficking of a focused cell, all combinations of 12 cell types other than the focused cell are candidates for the cell set to be placed as a reference. For the cell to be analyzed, differentially expressed genes (DEGs) were generated as a reference for all combinations, and the estimation performance of the Elastic Net was evaluated. In this evaluation, the samples in the dataset were those in which the target cells of the trafficking analysis showed fluctuation compared with the control group by flow cytometry.

First, the influence of all combinations of cells in the reference was evaluated using the correlation coefficient between the deconvolution-estimated value and the actual value measured by flow cytometry for neutrophils, monocytes, NK cells, and eosinophils (Fig. 3a). The correlations between the estimated and measured values when the top 10 and bottom 10 combinations were calculated, and differences in estimation in performance, were observed depending on the choice of reference cell sets (Figs 3b, c and S9). Furthermore, the optimized top 10 estimates outperformed the performance of existing methods such as FARDEEP, EPIC, CIBERSORTx, and DCQ [4,28–30], which were performed with LM13 as reference (Fig. S10). All cell name combinations including top 10 and bottom 10 used here are summarized in Supplementary File S1.

**Fig. 3.**
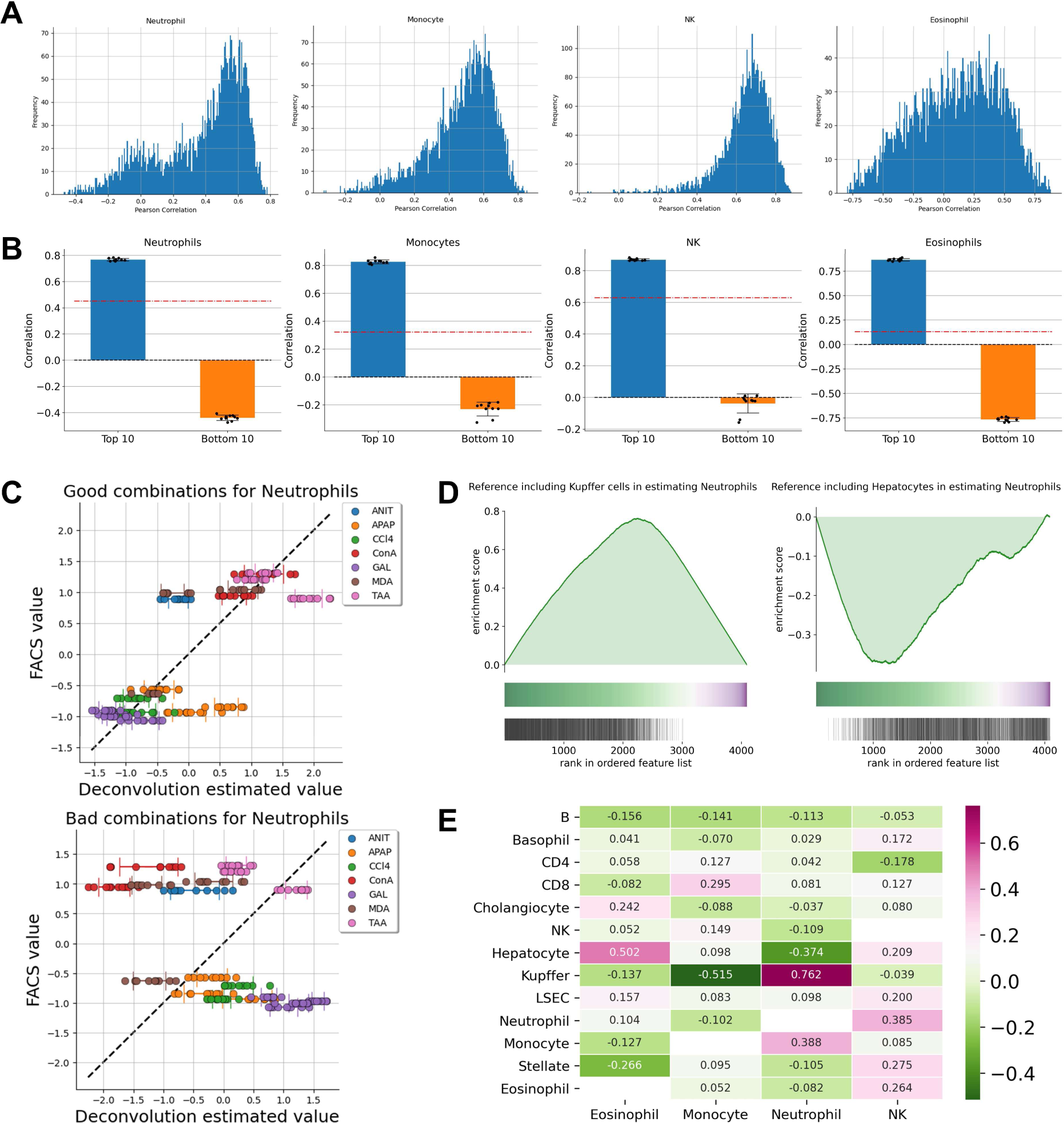
Optimization of reference cell type combinations. **A** Histogram showing the distribution of Pearson correlations when evaluating the performance of all possible cell type combinations as references. **B** Bar plots showing the difference between the top and bottom 10 Pearson correlations. The red line indicates the baseline when all 13 cell types are considered as reference. **C** Scatterplots showing the estimated and measured values with and without optimization. **D** Enrichment plot for the presence of Kupffer cells and hepatocytes in the reference for the estimation of neutrophils. The colored band represents the degree of correlation when estimated using each reference (green for a high correlation and purple for a low correlation). The bottom vertical black lines represent the location of the reference which include the Kupffer cells or hepatocytes. **E** Heatmap showing enrichment score when each cell is included in the estimation of the target cell.

Next, we evaluated the impact of the presence of a specific cell type in the reference cell set on the estimation performance of the cells under analysis. Specifically, when estimating neutrophil trafficking, combinations containing Kupffer cells tended to be enriched at the top with a high correlation coefficient, whereas combinations containing hepatocytes were enriched at the bottom with a low correlation coefficient (Figs 3d and S11a). Remarkably, the contribution of Kupffer cells and hepatocytes in the estimation of neutrophils was similar in the analysis using scRNA-Seq based reference (Fig. S11b). The combination of cells to be placed in the reference has good and bad affinity, and this also existed in the estimation of other cells (Fig. 3e). Therefore, we selected neutrophils, monocytes, NK cells, and eosinophils as the target cells for estimation due to computational cost issues and examined combinations of reference cells to establish a liver-specific optimized deconvolution model (Figs S12–S15).

The efficacy of the acquired reference cell set was appraised employing a pseudo-bulk dataset. Notably, in the estimation of neutrophils and NK cells, the utilization of the top 10 reference combinations exhibited superior estimation performance compared to the bottom 10 (see Figure S16). Conversely, it is important to note that the pseudo-bulk dataset retained linearity, resulting in marginal enhancements with the reference combination. These findings imply a potential disparity between the model optimized with real-world data and the artificial pseudo-dataset.

### Application to external public data

Evaluating extrapolation to aggregate immune cell findings from legacy data based on this model is essential. We then applied our optimized models based on reference cell combinations to external data for a more realistic task and evaluated their extrapolation and robustness. Bird et al. published transcriptome data at different time points after APAP administration (GSE111828) [31]. Because no immune cell information accompanies this dataset, we applied deconvolution using Elastic Net to each time point and evaluated the differences in performance with and without liver-specific optimization.

We compared the number of cell types constituting the top 10 and bottom 10 references to estimate the performance of the four cell types investigated for all reference cell combinations: neutrophils, monocytes, NK cells, and eosinophils (Fig. 4a). In preparing the references according to whether they are optimized, we attempted to eliminate the confounding effect of the number of cell types placed in the references on estimation performance. The threshold was set as the least number of cell types among the top 10 reference cell combinations found in the optimization process using the evaluation dataset. Nonoptimized combinations were selected for the bottom 10 combinations that retained cell types above the threshold. We performed deconvolution on GSE111828 with and without optimization, and the estimated values were also compared as a baseline when all 13 cell types were used. The results showed differences in the estimated percentage of immune cells at each time point after APAP administration (Fig. 4b). For example, the optimized model estimated an increase in eosinophils with a peak at 48 h after APAP administration, whereas the nonoptimized model had the opposite estimate of a decrease. The neutrophil estimates at 48 h postdose also showed opposite behaviors of increase and decrease, respectively, with and without optimization.

**Fig. 4.**
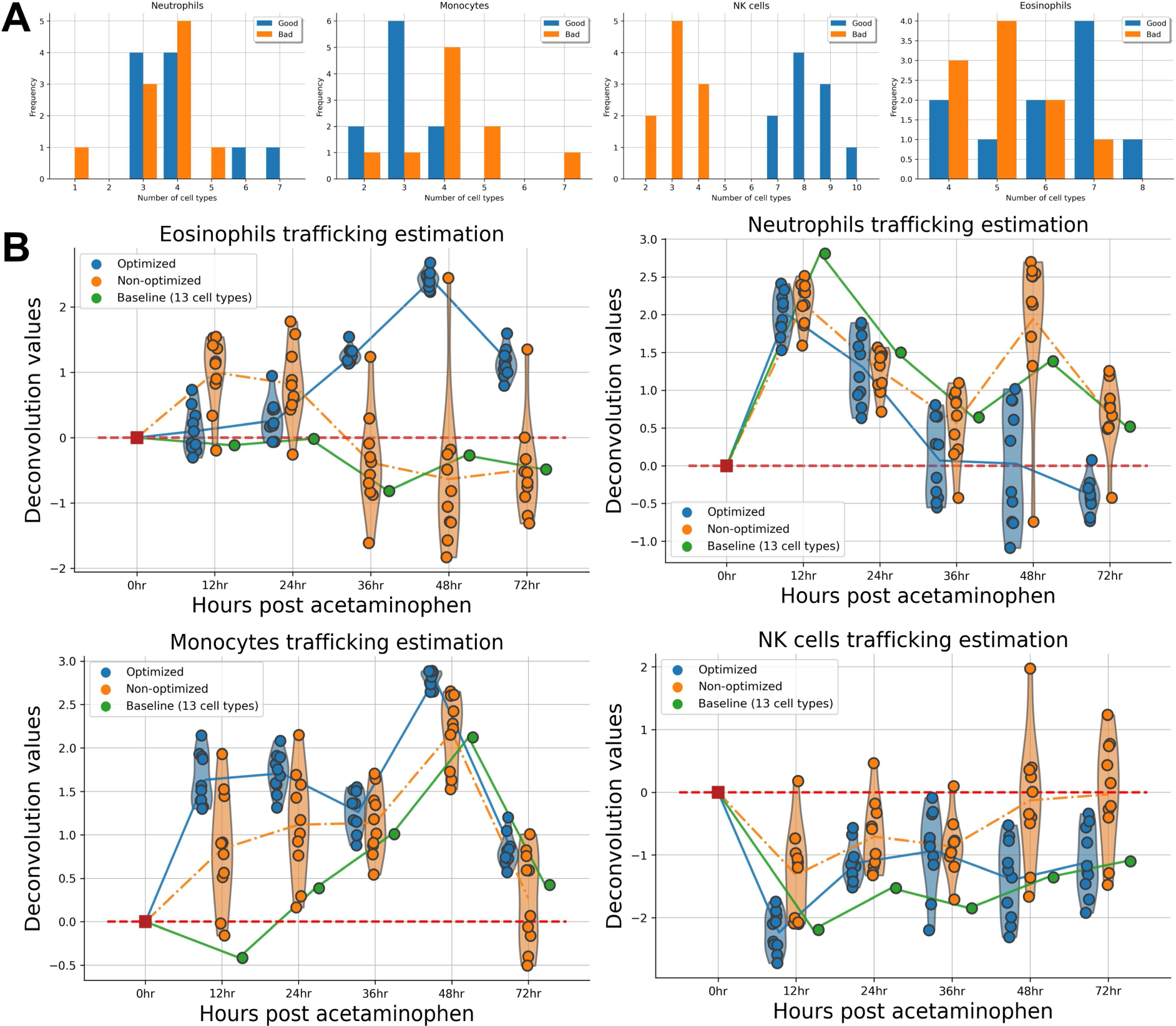
Application to external public liver tissue transcriptome data (GSE111828). **A** Bar plots showing difference of the number of cell types in top 10 and bottom 10 references in the estimation performance with the evaluation dataset. Blue bars and orange bars represent top 10 and bottom 10, respectively. **B** Estimated values when deconvolution was performed with and without liver tissue-specific optimization for each time point after APAP administration published in GSE111828. The violin plots at each time point represents the estimated values using top 10 (optimized) and bottom 10 (nonoptimized) references after eliminating confounding of the number of cells constituting the reference. The green line indicates the baseline when all 13 cell types are considered as reference.

Next, we evaluated the differences in the GSE111828 immune cell proportion estimates at each time point with and without the optimization described above to determine which reflected true immune cell behavior. We performed a reproduction study of APAP administration following the report of Bird et al. and measured immune cell proportions by flow cytometry to evaluate the extrapolation of the optimized model (Fig. 5a).

**Fig. 5.**
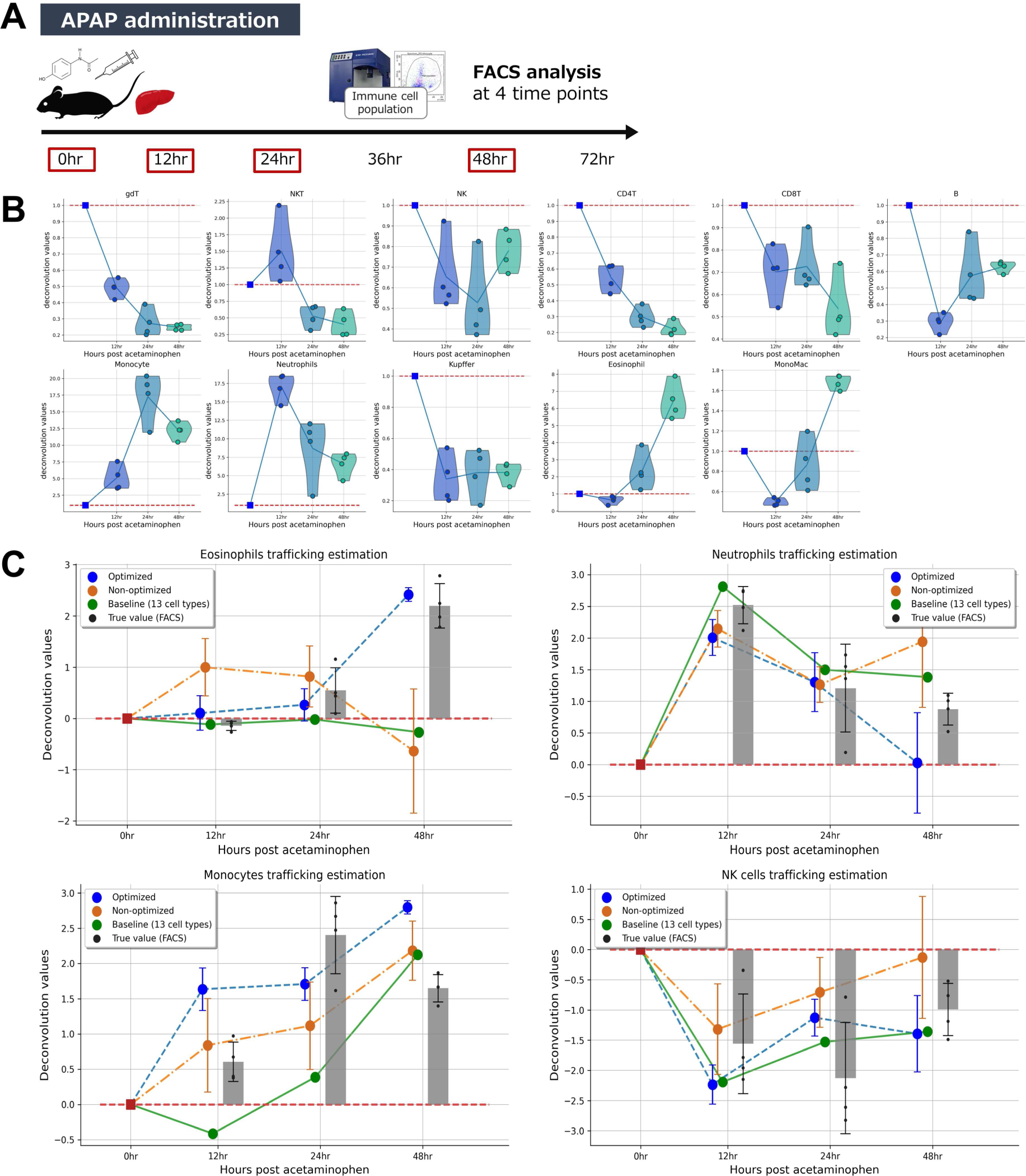
Reproduction of the APAP administration study and evaluating the effectiveness of the optimized model. **A** APAP was administered by single i.p. injection. Mice were euthanized 12, 24, and 48 h after APAP administration, and perfused liver samples were collected and subjected to flow cytometry analysis. **B** Violin plots showing the change in each immune cell proportion at each time point. The red line indicates the immune cell proportion of normal liver (0 h after administration). **C** Evaluation of the correspondence between the estimated values with and without liver-specific optimization and the measured values by flow cytometry. The blue and orange plot points indicate the mean estimated values by optimized and nonoptimized models, respectively. The green line indicates the baseline when all 13 cell types are considered as reference. Gray bar plots show the immune cell proportion measured by flow cytometry in this reproduction study. Error bars indicate the standard deviation.

For lymphocytes, six subtypes were selected as immune cells to be measured using flow cytometry: CD4+ T cells (CD4), CD8+ T cells (CD8), gamma-delta T cells (γδT), B cells, natural killer T (NKT) cells, and natural killer (NK) cells. For bone marrow-derived cells, measured immune cell proportion data were obtained for five subsets: monocytes, neutrophils, eosinophils, Kupffer cells, and monocyte-derived macrophage (MonoMac). Gating in flow cytometry of lymphocytes and bone marrow-derived cells is described in Figure S13. Each immune cell proportion at 12, 24, and 48 h after APAP administration was obtained and showed characteristic changes (Fig. 5b).

The correspondence between the estimated values and the actual values measured by flow cytometry for neutrophils, monocytes, NK cells, and eosinophils at each time point of GSE111828 was evaluated (Fig. 5c). Flow cytometry measurements showed an increase in eosinophils and a decrease in neutrophils at 48 h after APAP administration. These results were consistent with those estimated using the optimized reference, and establishing a tissue-specific deconvolution model enabled precise prediction. Furthermore, the findings are consistent with recent reports [32]. These results indicate that the combination of references optimized in this study is not merely overfitting to the prepared evaluation dataset but is highly extrapolative.

**Table. 1** Cell-specific gene expression profiles for 9 leukocyte subsets and 4 liver-related cell types were downloaded from the Gene Expression Omnibus (GEO). Hierarchical clustering with the Pearson correlation coefficient was performed on TPM-normalized profiles, and samples forming the main cluster were manually selected.

## Discussion

We established an evaluation dataset using various mouse liver injury models with corresponding transcriptome and immune proportions to evaluate the impact of modeling tissue-specific deconvolution. The small compounds used in the present study to induce diverse liver injuries contain unrecognized effects and provide novel findings that contribute to the understanding of immune cell trafficking. For example, 4,4′-methylene dianiline (MDA) and galactosamine (GAL) are mainly used for analysis as hepatotoxic substances in rats [29, 30], and there are few examples of comprehensive measurement of immune cell trafficking using flow cytometry in mice. In particular, the increase in eosinophils in liver tissue after GAL administration in this study is, to our knowledge, the first report providing new insights into the role of eosinophils in acute liver injury. Xu et al. reported that eosinophils play a protective role in acute liver injury induced by acetaminophen (APAP), CCL4, and ConA by accumulating at the site of injury [32]. In the present study, we observed an increase in eosinophils after CCL4 and ConA administration, supporting the results of Long Xu et al. and suggesting a similar protective role for GAL administration. The present study also revealed a tendency for eosinophils not to be induced in severely injured samples with high levels of alanine aminotransferase (ALT) after APAP administration. There is no stratification of eosinophil trafficking according to the degree of liver injury after APAP administration, and it is possible that the molecular mechanisms of hepatic protective function responsible for eosinophil trafficking may be disrupted during severe injury. These characteristic observations on eosinophils would contribute to the elucidation of a common molecular mechanism in the hepato-protective function of eosinophils in a variety of liver dysfunctions with different toxic action points and mechanisms.

In the optimization process of the reference cell sets using the established evaluation dataset, there was a correspondence between the biological meaning and the combinations. Kupffer cells are frequently included in combinations suitable for neutrophil estimation. This may be due to the multicollinearity of the deconvolution model. Although the Elastic Net employed in the deconvolution model of this study can handle multicollinearity with the L2 penalty to some degree [35], the effect itself is present. CD4+ T cells and CD8+ T cells showed similar gene expression profiles and were highly multicollinear (Fig. S5). Therefore, even if one is not included in the reference, the effect on the estimation performance of other cells, such as neutrophils, is small. In contrast, cell types with relatively low multicollinearity profiles, such as Kupffer cells, are considered to have a large effect on the estimated value depending on whether they are included in the reference or not. One reason why the inclusion or exclusion of Kupffer cells in the estimation of neutrophils and monocytes has a large impact on performance may be that all these cells are subtypes of myeloid cells. When performing reference-based deconvolution, it is important to consider variable combinations for variables derived from similar cell types with low multicollinearity.

The APAP administration study was conducted using the same design as that reported by Bird et al., and changes in immune cell proportions were measured by flow cytometry [31]. Neutrophils and monocytes peaked at 12 and 24 h after administration, respectively, whereas eosinophils showed a monotonous increase until the 48-h point. These results suggest that immune cells, such as neutrophils, monocytes, and eosinophils, may migrate to inflammatory sites differently after APAP administration. When estimating cell-specific immune trafficking using deconvolution, the optimized model showed outstanding estimation performance for eosinophils and neutrophils. These results strongly support the importance of tissue-specific and cell-specific deconvolution modeling and indicate that optimization within this study is highly extrapolative. Recent improvements in transcriptome analysis technology have made mouse liver data abundantly available in public databases. Although there are very few examples of observing the transition of the ratio of immune cells in the liver along a time axis, time-series transcriptome data do exist, as in the study by Bird et al. Applying the model optimized in the present study to these vast amounts of data is expected to enable us to obtain aggregated knowledge on immune cell trafficking in the liver.

The present study has several potential limitations. We took the liver as an example of a tissue and showed the optimization of tissue-specific deconvolution and its usefulness. The liver is a single organ with various disorders and is suitable in that it is possible to construct models with less confounding using small molecular compounds [36]. In addition, liver injury is a fundamental pathology of cirrhosis and subsequent liver cancer and the biggest cause of drug development discontinuation and market withdrawal, so understanding the behavior of immune cells has a large social impact [37,38]. On the other hand, when performing deconvolution for tissues other than the liver, it is necessary to prepare disorder models that reflect various immune cell trafficking to the target tissue and to construct an optimized model using the evaluation dataset. Additionally, for human clinical tissue samples, it is expected to be difficult to establish an evaluation dataset using compound perturbations, and the pipeline in the present study cannot simply be applied. Therefore, it is important to apply deconvolution methods other than reference-based methods that require tissue-specific optimization.

## Conclusions

In recent years, tissue transcriptome data have been abundantly accumulated in databases, and acquiring novel biological insights from these legacy data is an important and challenging task. Deconvolution methods that extract immune cell information from tissue transcriptome data are attractive options. Here, we established an evaluation dataset of the corresponding transcriptome and immune cell proportions in various mouse liver injury models using small compounds for liver-specific modeling. Using this dataset, we provided a methodology to optimize the combination of reference cell types for each cell to be estimated. The optimized model was applied to external data, suggesting that the model more accurately captures the time-series changes in each immune cell after APAP administration and is highly extrapolatable. The results presented here emphasize the importance of tissue-specific modeling when applying reference-based deconvolution methods to tissue transcriptome data.

## Methods

### Animals

Five-week-old and nine-week-old male C57BL/6JJcl mice were purchased from CLEA (Tokyo, Japan) and kept under standard conditions with a 12-hour day/night cycle and access to food and water ad libitum.

### Drug-induced liver injury models

Six-week-old mice acclimatized for one week were used for the experiments. Seven compounds, alpha-naphthyl isothiocyanate (ANIT, I0190, TCI, Japan), acetaminophen (APAP, H0190, TCI), CCl_4_ (CCl4, 039-01276, Fujifilm Wako, Japan), concanavalin A (ConA, 09446-94, Nacalai Tesque, Japan), galactosamine (GAL, G0007, TCI), 4,4′-methylene dianiline (MDA, M0220, TCI), and thioacetamide (TAA, T0187, TCI), were administered after 12 h of fasting. Depending on the compound to be administered, the vehicle was selected from 0.5% methylcellulose solution (133-17815, Fujifilm Wako), saline, and corn oil (032-17016, Fujifilm Wako). The administration route and concentration of each compound are listed in Supplementary Table S1. As a control group, 0.5% methylcellulose solution was used for oral administration, and saline was used for tail vein and intraperitoneal administration. The dose was 10 μL/kg for all groups. We euthanized animals 24 h after administration, and perfused liver samples and blood were harvested (*Liver and blood sample collection* section). For the obtained liver, a portion of the outer left lateral lobe of the liver was subjected to RNA isolation (*RNA-Seq analysis* section), and the remaining tissue was subjected to flow cytometry analysis (*Flow cytometry analysis* section).

### Time-dependent model of APAP-induced liver injury

Bird et al. performed RNA-Seq analysis on mouse liver tissue samples at different time points after APAP administration [31], and we followed this protocol.

Ten-week-old mice acclimatized for one week were fasted for 10 h from 7 am to 5 pm. APAP was dissolved in sterile phosphate-buffered saline (PBS) warmed to 42°C and administered at 350 mg/kg by a single i.p. injection of 20 μL/g. Animals were sacrificed 12, 24, and 48 h after APAP administration, and perfused liver samples and blood were collected (*Liver and blood sample collection* section). The harvested liver sample was subjected to flow cytometry analysis (*Flow cytometry analysis* section).

### Liver and blood sample collection

Briefly, a superior vena cava was clipped using a clamp, and the blood was collected through an inferior vena cava into a 1.5-mL tube containing 1 μL heparin (Yoshindo, Toyama, Japan). The collected blood sample was centrifuged (800×*g*, 4°C for 15 min) for serum separation. Serum alanine aminotransferase (ALT), aspartate aminotransferase (AST), and total bilirubin (TBIL) were measured using a Dri-Chem NX500sV (Fujifilm Corporation). After cutting the portal vein, the first perfusion was performed by injecting 10 mL of 5 mM HEPES (H4034, Sigma-Aldrich, USA)/5 mM EDTA (345-01865, Fujifilm Corporation) Hanks’ Balanced Salt Solution (17461-05, Nacalai Tesque) through the inferior vena cava. Then, the second perfusion was performed by using 10 mL of 5 mM HEPES Hanks’ balanced salt solution. Before harvesting the tissue, 2 mL of dissociation enzyme solution of gentleMACS (Miltenyi Biotec, Germany) was filled in the liver from an inferior vena cava by clipping the cut portal vein.

### RNA-seq analysis

Total RNA was prepared using Isogen II (311-07361, Nippon Gene, Japan) and purified using an RNeasy Plus Mini Kit (74136, Qiagen, Netherlands) with gDNA elimination by an RNase-Free DNase Set (79254, Qiagen), following the manufacturer’s protocols. RNA-Seq libraries were prepared with a TruSeq Stranded mRNA Sample Preparation kit (Illumina, USA). The libraries were sequenced for single-end reading using a NovaSeq 6000 (Illumina).

Quality control of all reads was performed using PRINSEQ++ (version 1.2.4) with the indicated parameters (trim_left = 5, trim_tail_right = 5, trim_qual_right = 30, ns_max_n = 20, min_len = 30) [39]. The expression of transcripts was quantified using Salmon (version 1.6.0) and gencode.vM28.transcripts obtained from GENCODE with the indicated parameters (validation Mappings, gcBias, seqBias) and decoy-aware index created using Salmon and GRCm39.primary_assembly.genome obtained from GENCODE [40,41]. Transcripts per kilobase million (TPM) data were obtained using tximport, which is implemented in the software package Bioconductor with R (version 4.1.3) from quant.sh files created by Salmon.

### Flow cytometry analysis

The liver sample was dissociated using gentleMACS, according to the manufacturer’s instructions. Except where noted, PBS containing 2% fetal bovine serum was used as “wash buffer” thereafter. The washed samples were centrifuged (50 ×*g*, 4°C for 3 min) to eliminate hepatocytes and were subjected to ACK lysis buffer. ACK buffer was prepared by adding 8,024 mg of NH_4_Cl (A2037, TCI), 10 mg of NHCO_3_ (166-03275, Fujifilm Wako), and 3.772 mg of EDTA 2Na·2H_2_O (6381-92-6, Dojindo Laboratories, Japan) into 1 L of pure water. The samples were washed with wash buffer three times, and then the samples were subjected to flow cytometry analysis. Flow cytometric analysis was performed with FACSAria III (BD Biosciences, USA), and data were analyzed with FlowJo software (Treestar). Details are provided in the Supplementary Note.

### Deconvolution approach

#### Elastic Net

Consider a measured bulk gene expression matrix ***Y*** ∈ ***R^N×M^*** for N genes across M samples, each containing K different cell types. The goal of deconvolution is to estimate cell type-specific expression ***X*** ∈ ***R^N×K^*** and cell type proportion matrix ***P*** ∈ ***R^K×M^*** and can be written as:

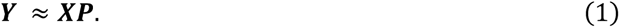

Elastic Net [35] is a regularized regression model with combined L1 and L2 penalties. We can estimate the cell type proportion matrix 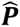 via:

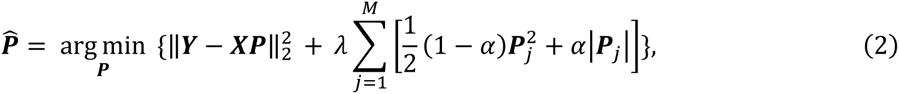

where λ and α are hyperparameters, and we set λ = 1 and α = 0.05 as default parameters.

### Bulk tissue gene expression matrix

The TPM-normalized liver expression profile is the analysis target. The transcript IDs were converted to MGI gene symbols using files available from Biomart [42], and median values were selected for duplicate gene names. The data were processed in the following order: log transformation, elimination of low-expressing genes, batch normalization, and quantile normalization. Among the genes in this normalized expression matrix, we focused on N marker genes, which are described below, and defined them as ***Y*** ∈ ***R^N×M^***. These preprocessing steps can be reproduced in our repository [43].

### Cell type-specific expression matrix (Reference)

#### Bulk RNA-Seq-derived reference

First, we downloaded raw RNA-Seq expression datasets for 9 leukocyte subsets and 4 liver-related cell types from the Gene Expression Omnibus (GEO). Hierarchical clustering with the Pearson correlation coefficient was performed on TPM-normalized profiles, and samples forming the main cluster were manually selected. Note that data are available at our GitHub repository [43]. Then, we converted transcript IDs to MGI gene symbols, and median values were selected for duplicate gene names. The data were processed in the following order: log transformation, elimination of low-expressing genes, and quantile normalization. Only genes in common with the bulk tissue gene expression matrix were retained.

#### Single cell RNA-Seq-derived reference

Single-cell RNA-Seq data of liver cells in a mouse model of acute liver failure was accessible on ArrayExpress under accession E-MTAB-8263 [DOI: 10.1038/s41591-020-1102-2]. Processed data was downloaded, resulting in the acquisition of 5 leukocyte subsets and 5 cell types relevant to the liver. These cell types align with those encompassed in the bulk RNA-Seq-derived reference.

#### Detection of differentially expressed genes (DEGs)

Gene expression profiles specific to each of the K cell types constituting the reference were selected as differentially expressed genes. We retained up to 50 genes as markers, exhibiting an absolute fold change surpassing 1.5 for the second cell type with the highest expression. Subsequently, N genes sourced from the aforementioned K-cell markers were encompassed in the analysis, leading to the definition of the cell type-specific expression matrix ***X*** ∈ ***R^N×K^***. For the bulk RNA-Seq-derived reference, a total of 503 DEGs spanning 13 cell types were identified, while the scRNA-Seq-derived reference yielded 184 DEGs across 10 cell types (Supplementary Files S2 and S3). The code for this detection is integrated into our proposed pipeline and is accessible in our GitHub repository [43].

### Evaluation

The estimated cell type proportion matrix 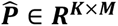 was obtained by running deconvolution and converted to a sample-wide *z* score for each cell of interest. Similarly, the ground truth cell type proportion matrix obtained by flow cytometry was converted to a sample-wide *z* score, and its Pearson correlation with 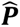 was evaluated. Note that the estimated 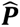 is not necessarily greater than 0 because it is not constrained to be nonnegative.

### Optimization of the combination of reference cell types for immune cell trafficking

#### Selection of samples showing fluctuation

We used our established evaluation dataset to optimize the best combination of reference cell types for estimating the trafficking of the immune cell of interest. In the optimization, neutrophils, monocytes, eosinophils, and NK cells were selected for trafficking compared with the control group, and samples showing fluctuations in cell proportions were evaluated. The amount of change for each sample was plotted, and samples outside the quartile range were considered to have a large fluctuation in immune cells and were used in the optimization. Note that samples that fall below Q1 − 1.5×IQR or above Q3 + 1.5×IQR were considered outliers and removed from optimization. Here, IQR is the interquartile range, and Q1 and Q3 are the lower and upper quartiles, respectively.

#### Optimized reference selection

Reference-based deconvolution can estimate the proportion of cells in the reference. Therefore, in the analysis of immune cell trafficking, the target cell must be included in the reference, and if K types of cells are candidates for the reference, there are a total of 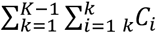 combinations. For each immune cell of interest, deconvolution was performed on all cell combinations comprising the reference for samples showing fluctuation. Pearson correlations between sample-wide transformed *z* scores were calculated for the deconvolution output and the measured values using flow cytometry. Among the Pearson correlation scores performed on all obtained references, the 10 highest references were considered to be the optimized reference.

### Evaluation of extrapolation to external data (GSE111828)

Bird et al. performed RNA-Seq analysis on mouse liver samples following acetaminophen-induced liver injury. Transcription profiles were generated at 12, 24, 36, 48, and 72 h after injury and were publicly available as GSE111828. Using the optimized combination of cell types in the reference, we estimated the changes in immune cell trafficking at each time point. Furthermore, we evaluated the estimation performance by reproducing the experiment under the same conditions and obtaining the ground truth of the changes in the immune cell ratio by flow cytometry (*Time-dependent model of APAP-induced liver injury* section).

### Pseudo-bulk dataset from scRNA-Seq

We engineered an artificial pseudo-bulk amalgamation by randomly extracting individual cells quantified through scRNA-Seq and aggregating them to constitute a total of 1000 cells. To ensure representation, we stipulated the inclusion of hepatocytes among the 10 cell types present. Subsequently, we designated the proportion assigned to each cell type. Notably, hepatocytes were allocated proportions ranging from 0.7 to 0.8, while a sum-to-one constraint was enforced across all cell types. This process was reiterated, resulting in the formulation of a pseudo-bulk dataset encompassing 1000 samples.

## Availability of Source Code and Requirements

Project name: LiverDeconv

Project home page: https://github.com/mizuno-group/LiverDeconv

Programming languages: Python 3.8

Other requirements: combat, numpy, matplotlib, pandas, scipy, sklearn, statsmodels

Operating systems: Linux, Windows

License: MIT

## Data Availability

Code, models, and data used in this article are available on GitHub page [43]. A total of 57 RNA-Seq data samples from various mouse liver injury models obtained in this study have been deposited with accession code GSE237801 in NCBI Gene Expression Omnibus. Processed RNA-Seq data and flow cytometry measurements for assessment are available in our GitHub repository, designed for convenient utilization.

## List of abbreviations

ALT: alanine transaminase;
ANIT: α-isothiocyanate;
APAP: acetaminophen;
ConA: concanavalin A;
DEGs: differentially expressed genes;
FACS: Fluorescence Activated Cell Sorter;
GAL: galactosamine;
GEO: Gene Expression Omnibus;
LR: Linear Regression;
MDA: 4,4’-methylene dianiline;
PCA: Principal Component Analysis;
sc RNA-Seq: single cell RNA-Seq;
TAA: thioacetamide.

## Declarations

### Consent for publication

Not applicable.

### Contributions

IA: Data curation, Formal analysis, Methodology, Software, Investigation, Writing – Original draft, Visualization.

TM: Conceptualization, Resources, Supervision, Project administration, Writing – Original draft, Writing – Review and editing, Funding acquisition.

KM: Data curation, Investigation.

YS: Methodology, Investigation.

HK: Writing – Review.

### Ethics approval and consent to participate

The studies reported in this article were performed in accordance with the guidelines provided by the Institutional Animal Care Committee (Graduate School of Pharmaceutical Sciences, the University of Tokyo, Tokyo, Japan).

## Acknowledgments

We greatly appreciate Dr. Bird’s advice on the protocol used to model APAP-induced liver injury in GSE111828. We thank all those who contributed to the construction of the following datasets employed in the present study, such as LM6, 9, and 13.

## Supplementary figures

**Figure S1.**
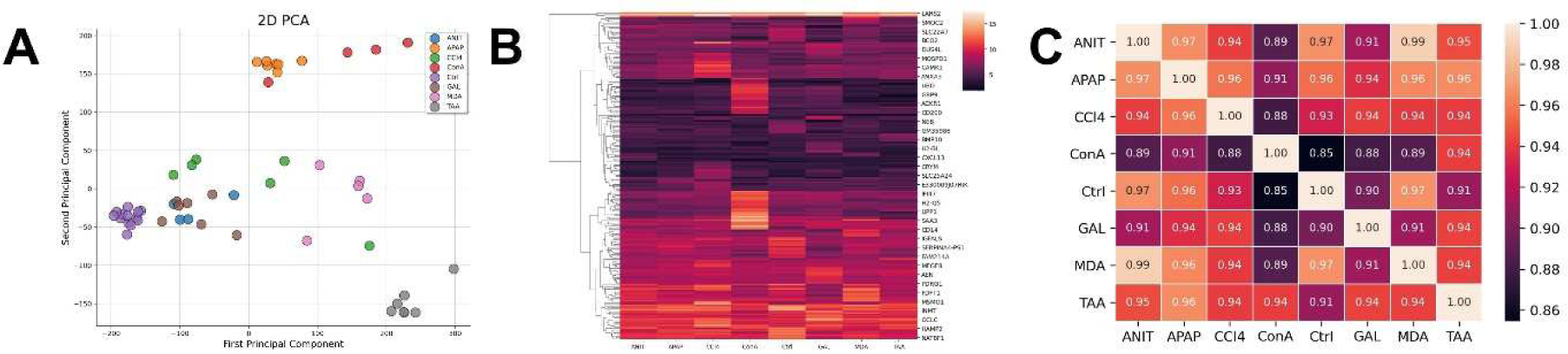
Gene expression profiles in diverse liver injury models. **A** PCA plotting (2D) of the obtained RNA-seq data shows clusters formed by each compound administration. **B** Heatmap showing mean gene expression level (*z* score) whole genes in each administration group. **C** Heatmap showing the Pearson correlation of expression profile of each administration group.

**Figure S2.**
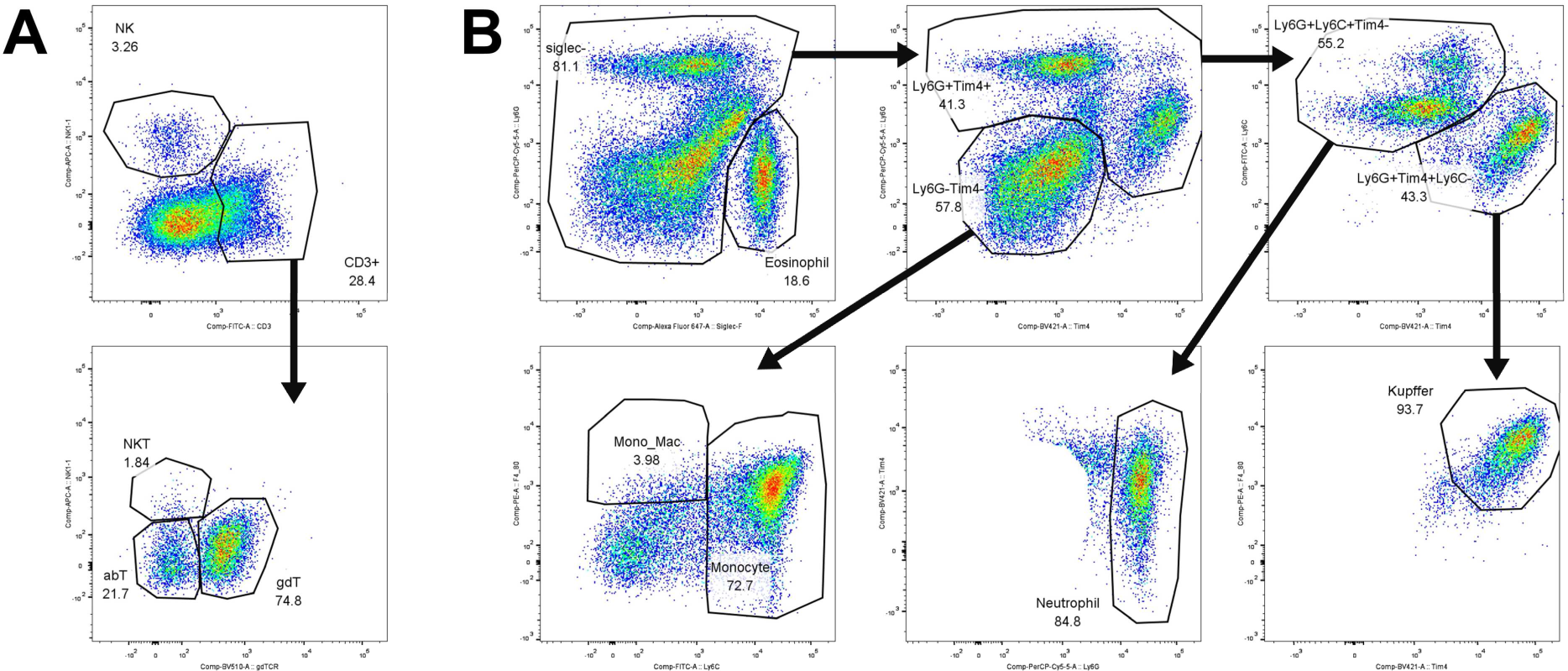
Flow cytometry gating strategy to establish an evaluation dataset using diverse mouse liver injury models. **A** Lymphocytes: αβT cells, γδT cells, natural killer T (NKT) cells, natural killer (NK) cells. **B** Myeloid cells: monocytes, neutrophils, Kupffer cells, eosinophils, and monocyte-derived macrophages (MDMs).

**Figure S3.**
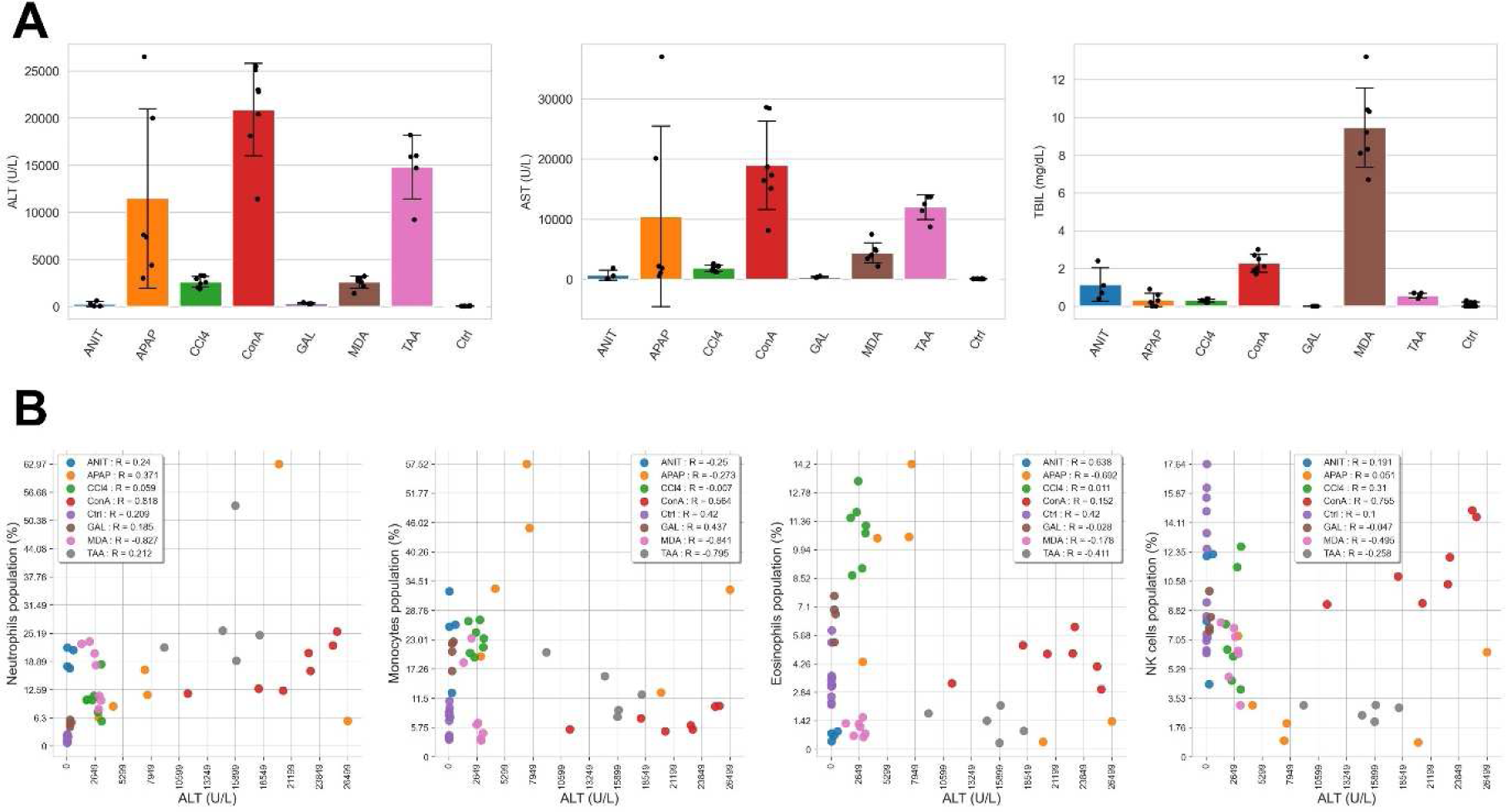
Blood biochemical values and immune response in each liver injury model. **A** Bar plots showing measurements of alanine aminotransferase (ALT), aspartate aminotransferase (AST), and total bilirubin (TBIL) in each administration group. Error bars represent the standard deviation of the mean. **B** Scatterplot showing the relationship between the accumulation of neutrophils, monocytes, eosinophils, and NK cells and the degree of liver injury. The higher the ALT value in the sample, the more severe the liver damage. The figures in legends represent the Pearson correlation within each group.

**Figure S4.**
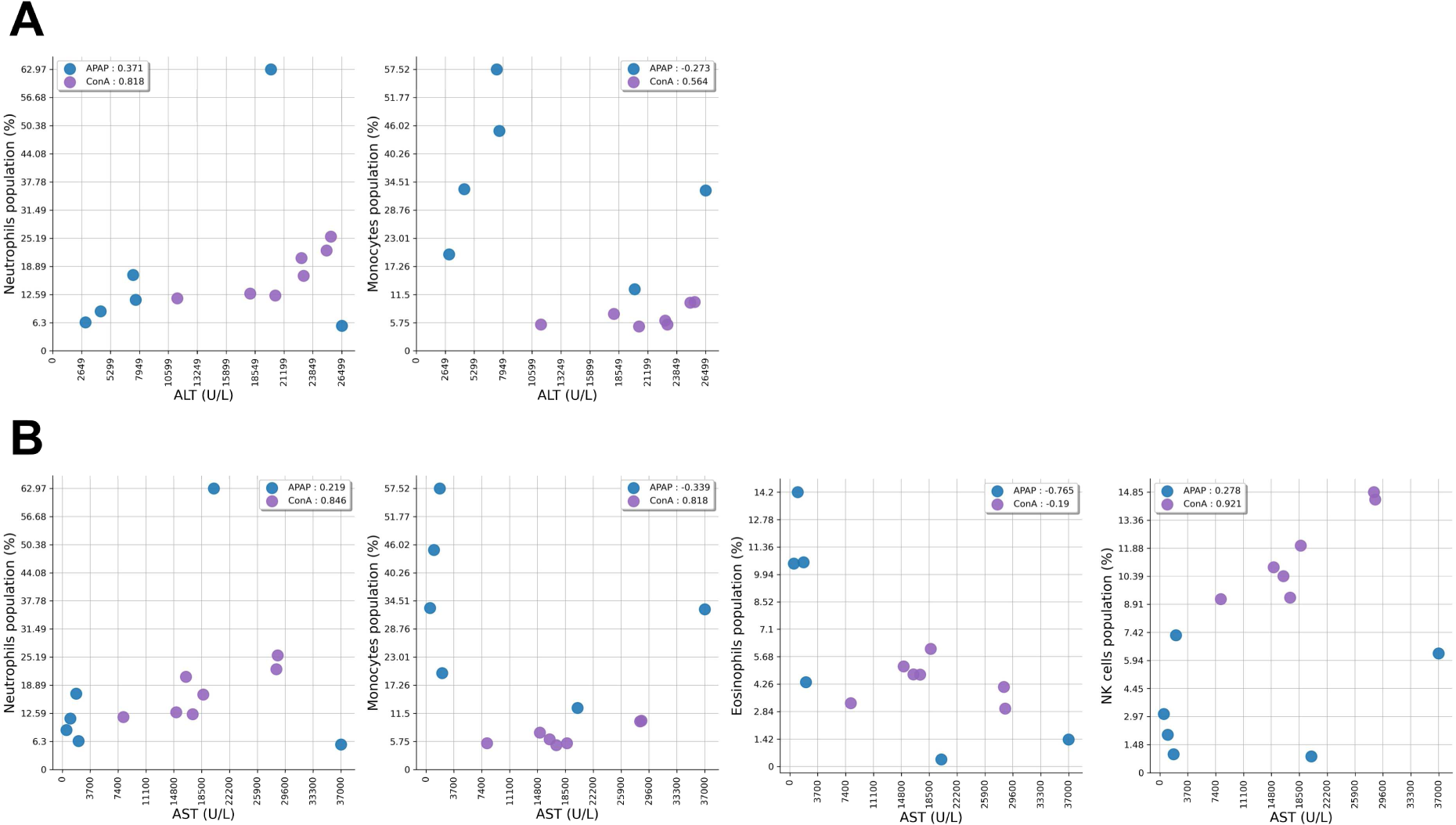
Relationship between the degree of liver injury and immune response after APAP and ConA administration. The figures in legends represent the Pearson correlation within each group. **A** The X axis indicates the alanine aminotransferase (ALT). **B** The X axis indicates the aspartate aminotransferase (AST).

**Figure S5.**
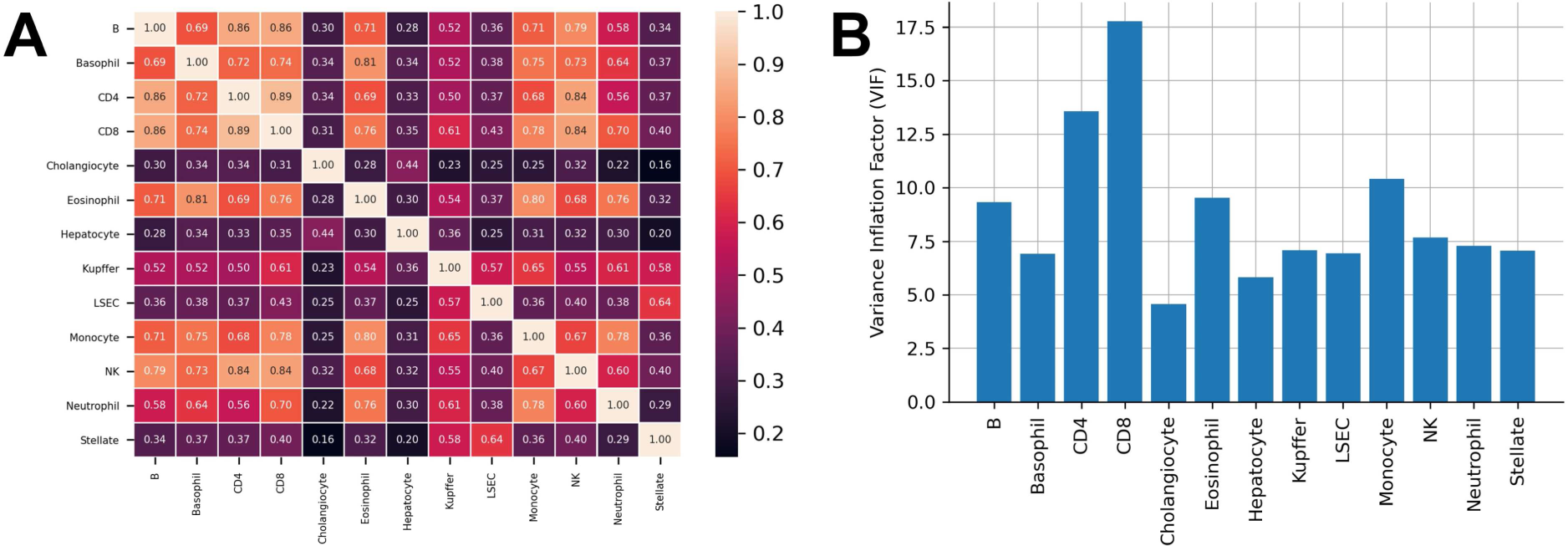
Similarity of differentially expressed gene profiles of the obtained 13 cell types. **A** Heatmap showing the Pearson correlation of median gene expression level of each cell type. **B** Bar plots showing the variance inflation factor value, an indicator for detecting multicollinearity.

**Figure S6.**
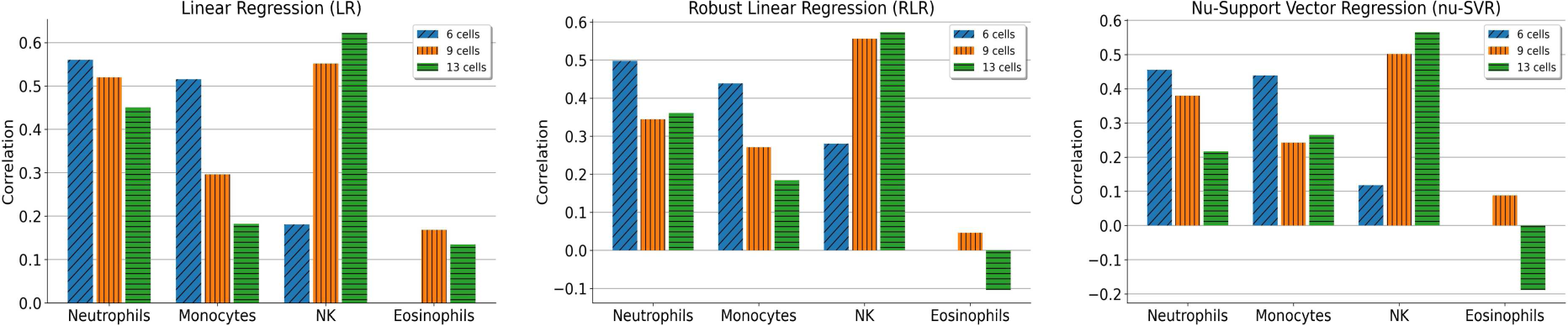
Effects of cell types constituting a reference on the estimation performance of representative reference-based deconvolution methods. Bar plots show the Pearson correlation when 6 cells (LM6), 9 cells (LM9), and 13 cells (LM13) are used as reference.

**Figure S7.**
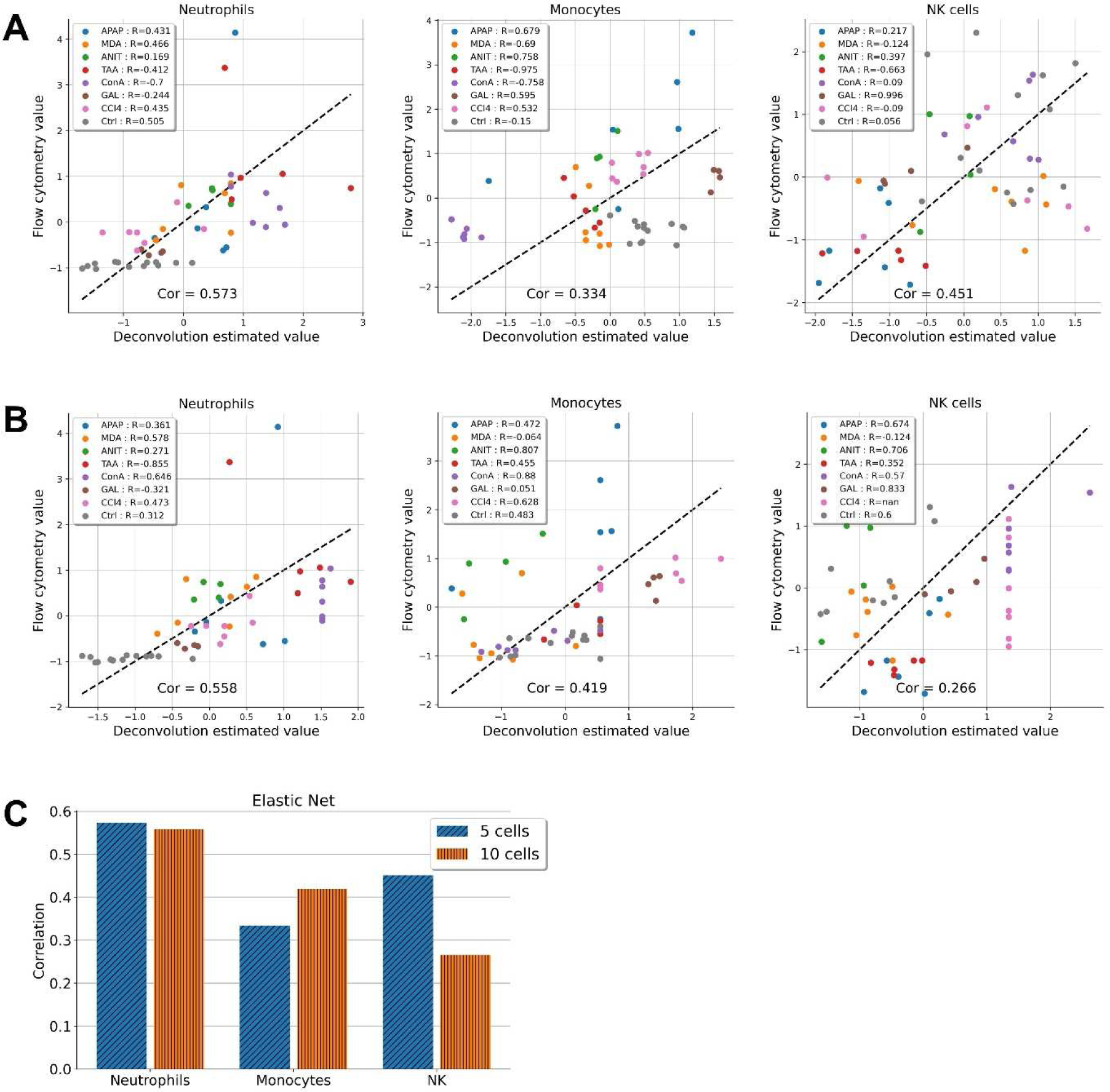
Conducting a deconvolution method on the dataset for evaluation by using scRNA-Seq as reference. The scatterplot shows the Pearson correlation between flow cytometry values and deconvolution estimates using Elastic Net when **A** 5 immune-related cell types and **B** 10 cell types are used as reference. **C** Bar plots showing the summary of the impact of the reference cell set on estimation performance.

**Figure S8.**
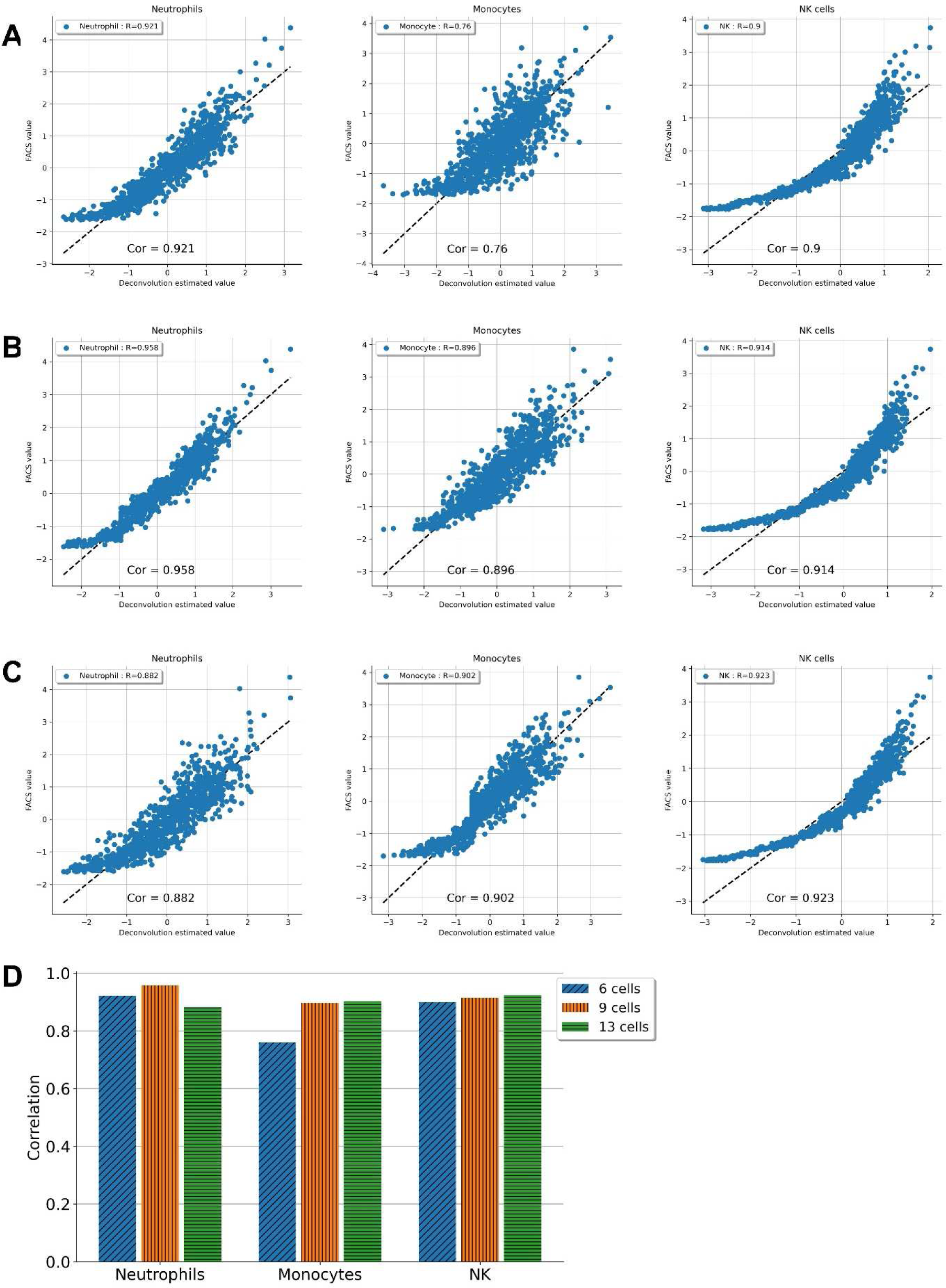
Conducting a reference-based deconvolution method on the artificial pseudo-bulk dataset. The scatter plot shows the Pearson correlation between flow cytometry values and deconvolution estimates using Elastic Net when **A** LM6, **B** LM9, and **C** LM13 are used as reference. **D** Bar plots showing the summary of the impact oof the reference cell set on estimation performance.

**Figure S9.**
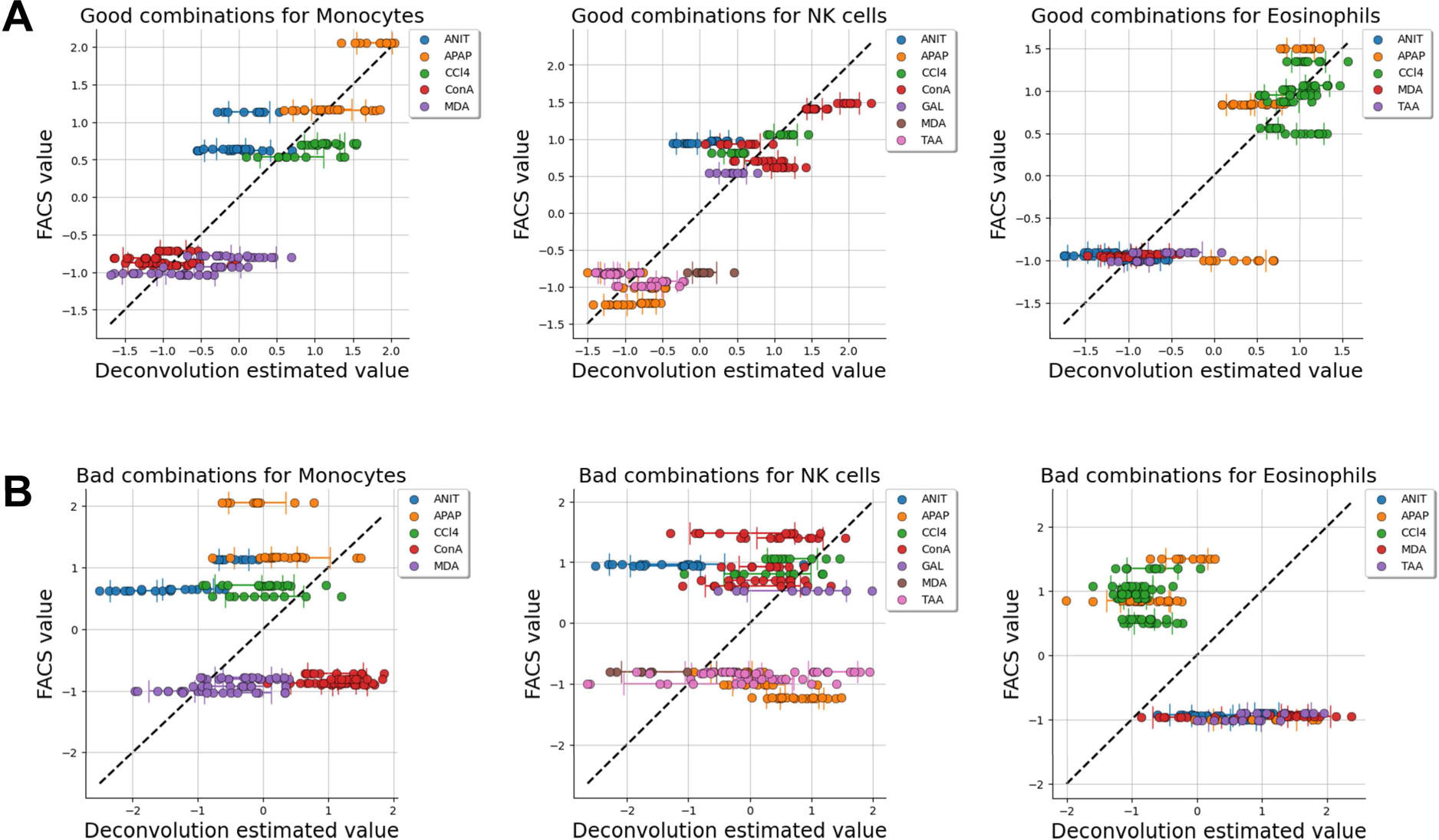
Scatterplots showing the deconvolution estimated values and true values measured by flow cytometry. **A** Estimation results using the 10 references with the best estimation performance using the evaluation dataset. **B** Estimation results using the 10 references with the worst estimation performance using the evaluation dataset.

**Figure S10.**
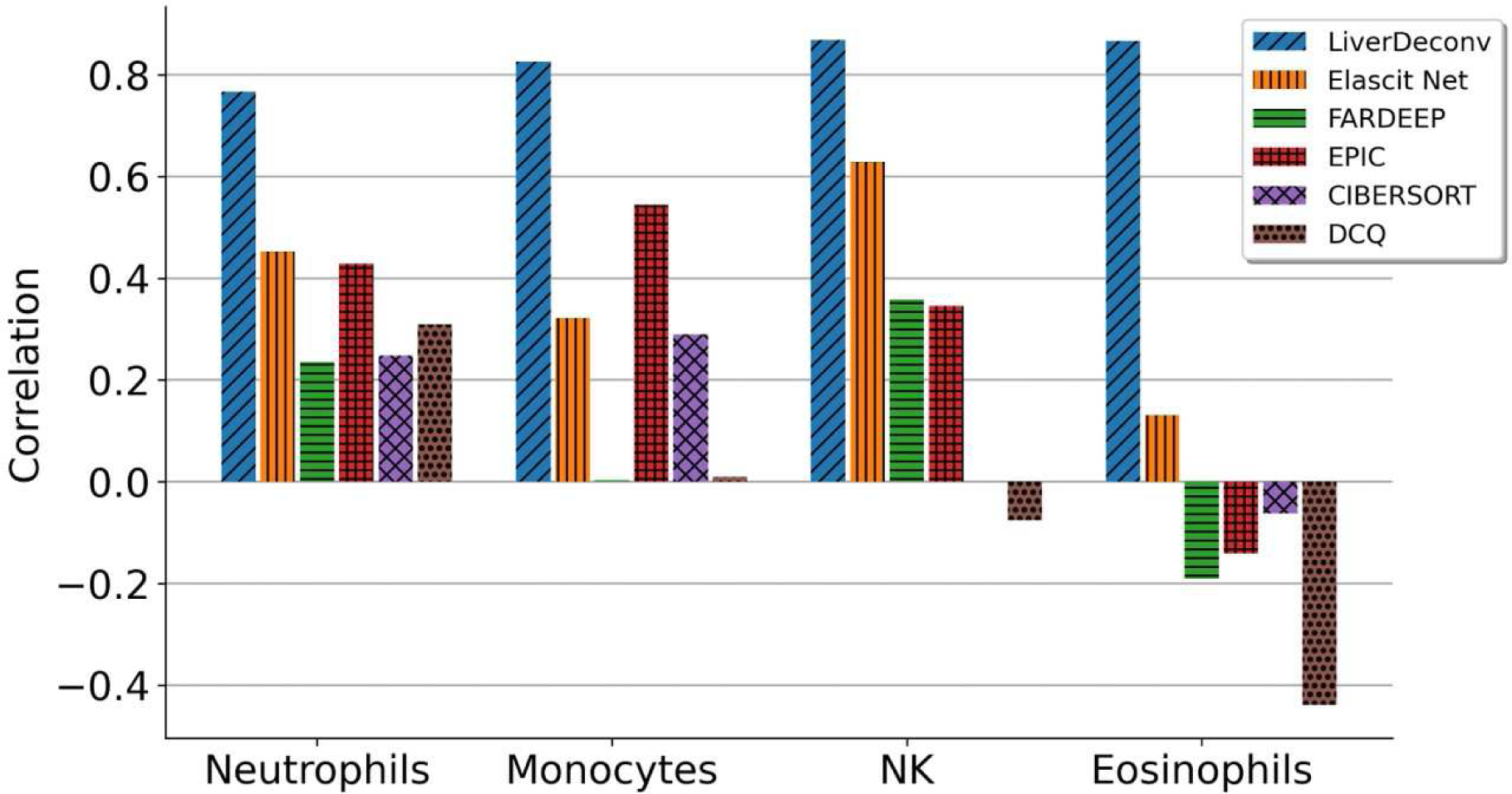
Bar plots showing the estimation performance of our optimized LiverDeconv method against currently available representative deconvolution tools such as FARDEEP, EPIC, CIBERSORTx, and DCQ.

**Figure S11.**
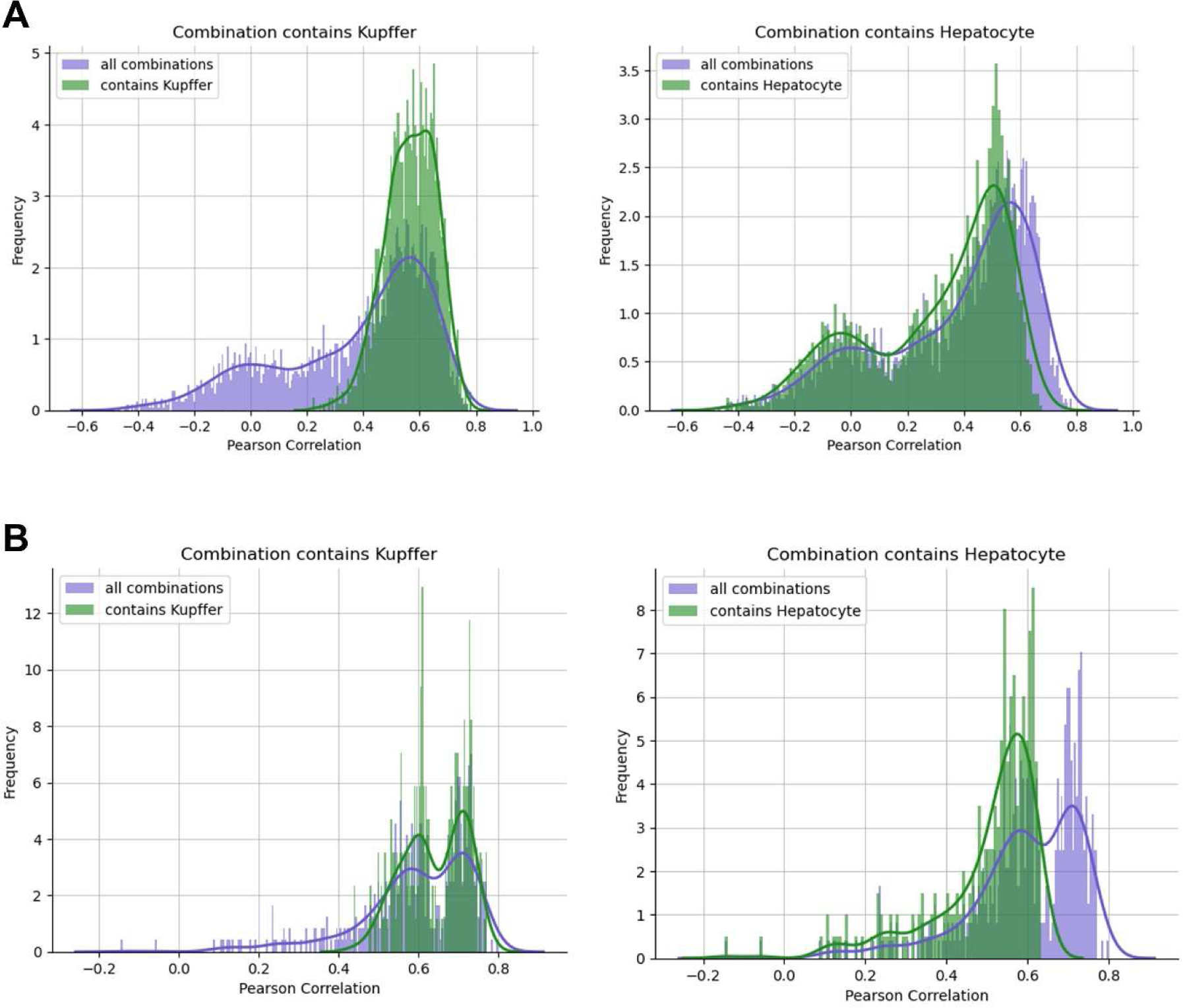
Bias in the distribution of correlation coefficients arises when specific cells are included in the reference for neutrophil estimation. The purple distribution represents all combinations of cell types, with the stipulation that neutrophils—the focus of analysis—are included. The green distribution further imposes the requirement that a specific cell type, such as Kupffer cells or hepatocytes, is included alongside neutrophils. **A.** Bulk RNA-Seq and **B.** scRNA-Seq were used as a reference, respectively.

**Figure S12.**
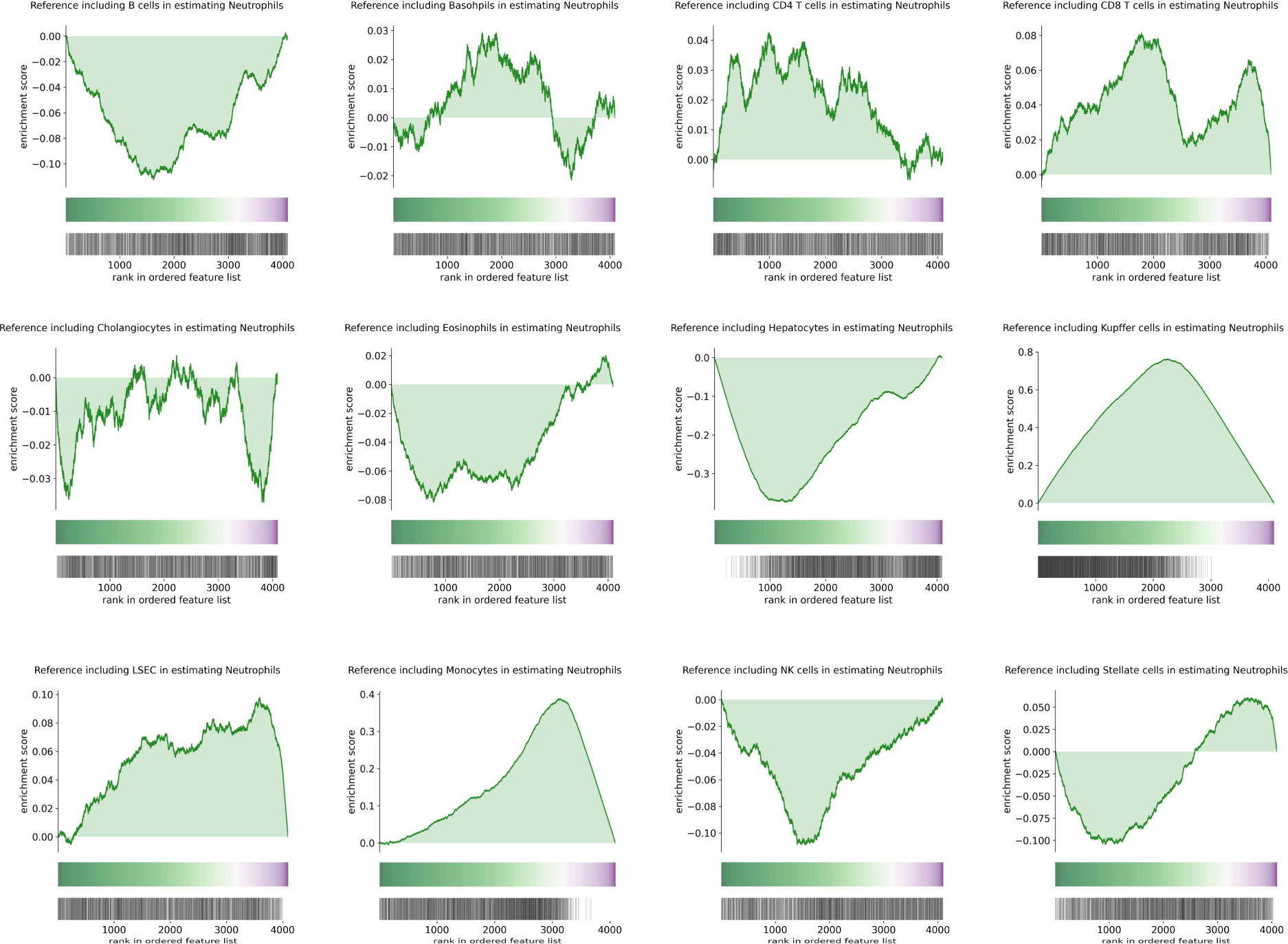
Evaluation of the effect on estimation performance of whether or not each other cell is included in the reference when estimating neutrophils. Enrichment plot for the presence of each cell in the reference. The Y axis indicates the enrichment score. The colored band represents the degree of correlation when estimated using each reference (green for a high correlation and purple for a low correlation). The bottom vertical black lines represent the location of the reference that includes the target cell.

**Figure S13.**
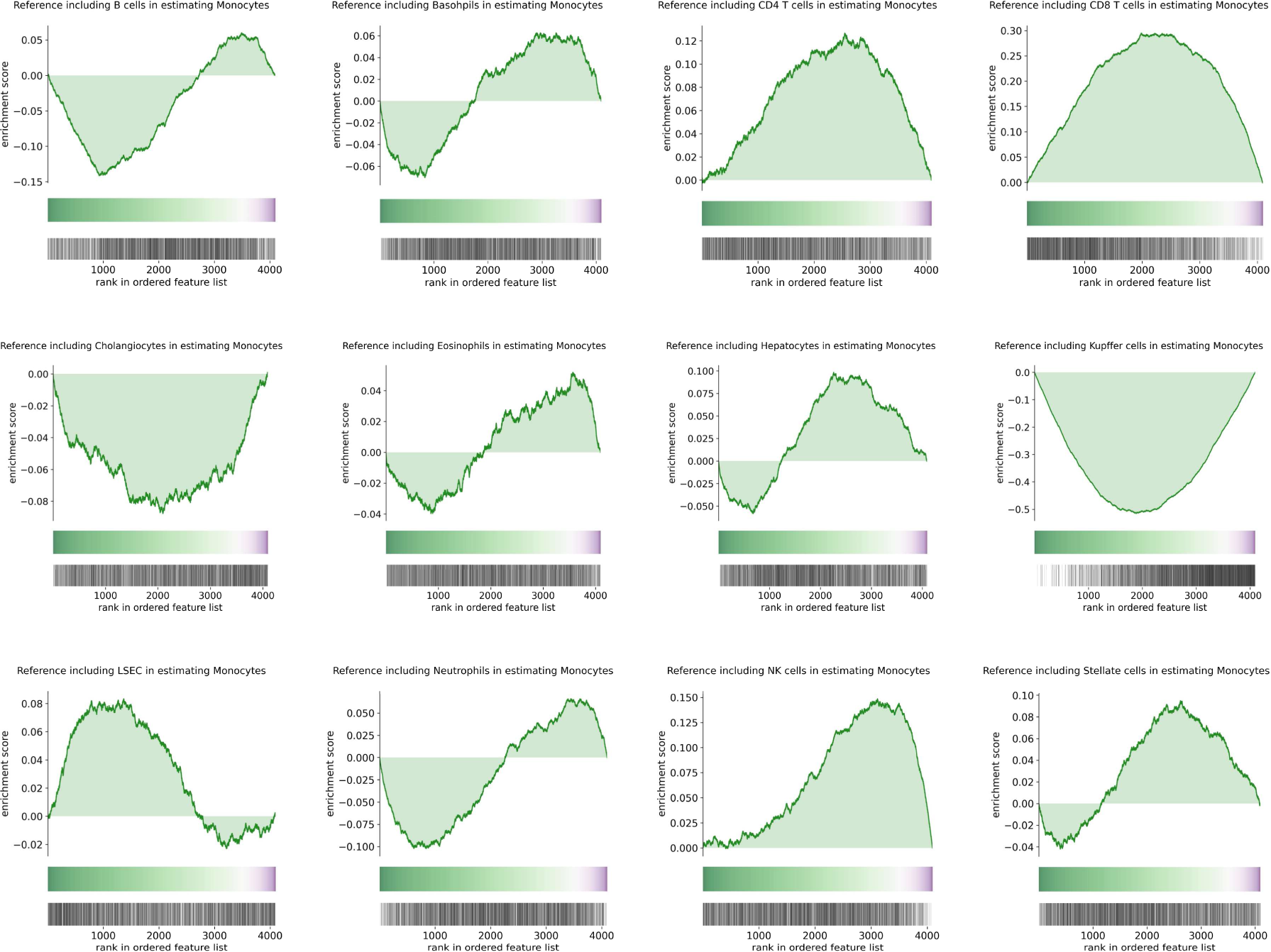
Evaluation of the effect on estimation performance of whether each other cell is included in the reference when estimating monocytes. Enrichment plot for the presence of each cell in the reference. The Y axis indicates the enrichment score. The colored band represents the degree of correlation when estimated using each reference (green for a high correlation and purple for a low correlation). The bottom vertical black lines represent the location of the reference that includes the target cell.

**Figure S14.**
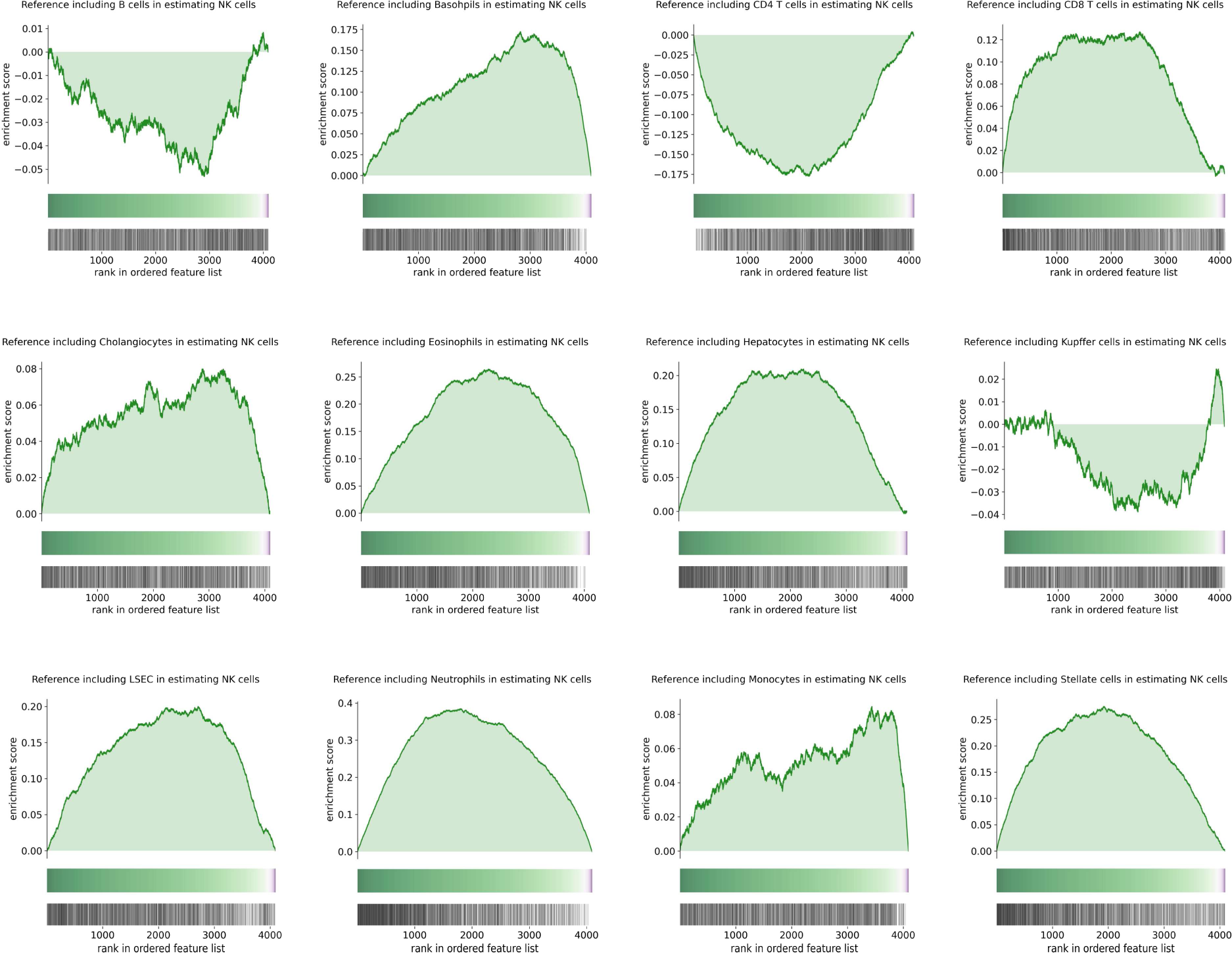
Evaluation of the effect on estimation performance of whether or not each other cell is included in the reference when estimating NK cells. Enrichment plot for the presence of each cell in the reference. The Y axis indicates the enrichment score. The colored band represents the degree of correlation when estimated using each reference (green for a high correlation and purple for a low correlation). The bottom vertical black lines represent the location of the reference that includes the target cell.

**Figure S15.**
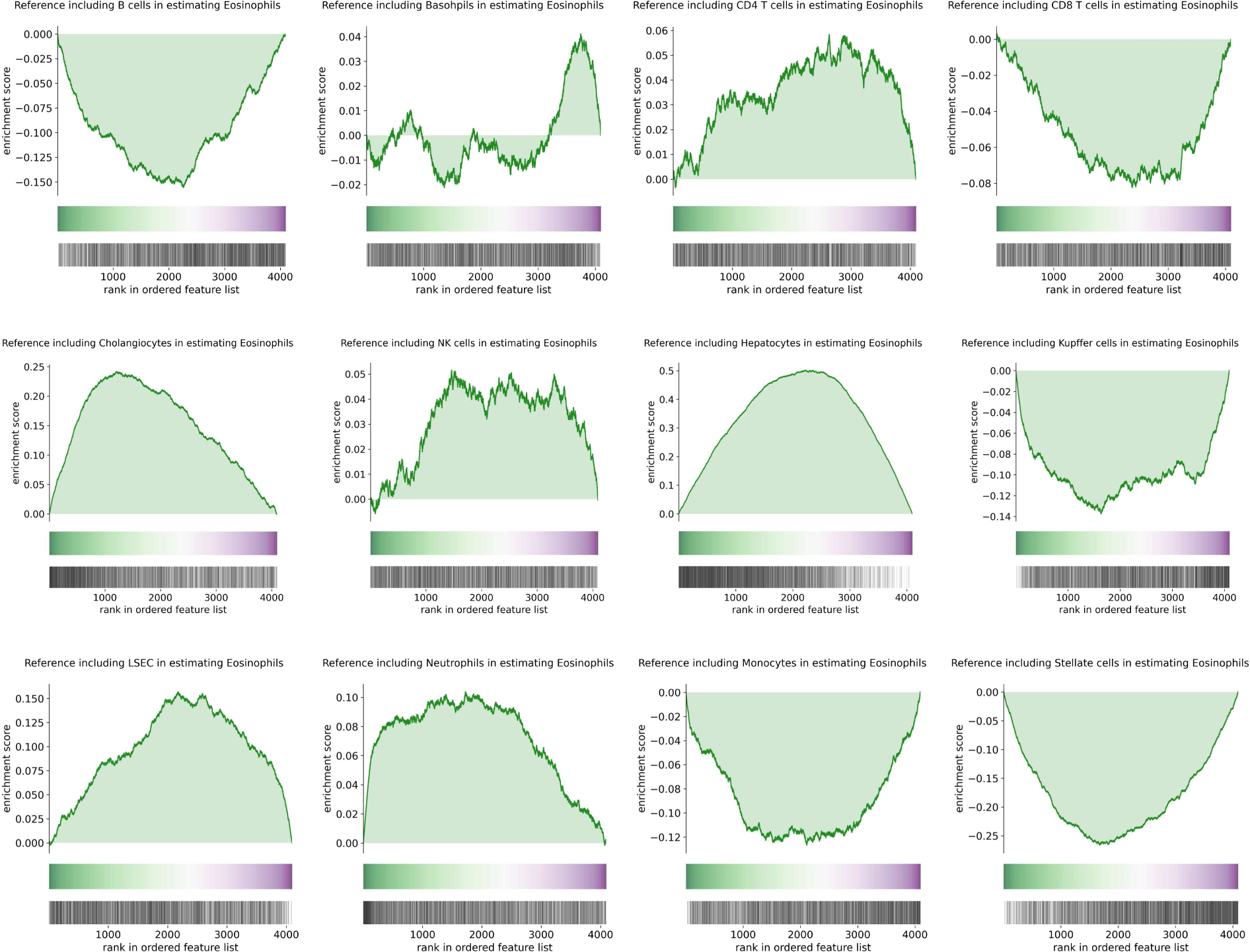
Evaluation of the effect on estimation performance of whether or not each other cell is included in the reference when estimating eosinophils. Enrichment plot for the presence of each cell in the reference. The Y axis indicates the enrichment score. The colored band represents the degree of correlation when estimated using each reference (green for a high correlation and purple for a low correlation). The bottom vertical black lines represent the location of the reference that includes the target cell.

**Figure S16.**
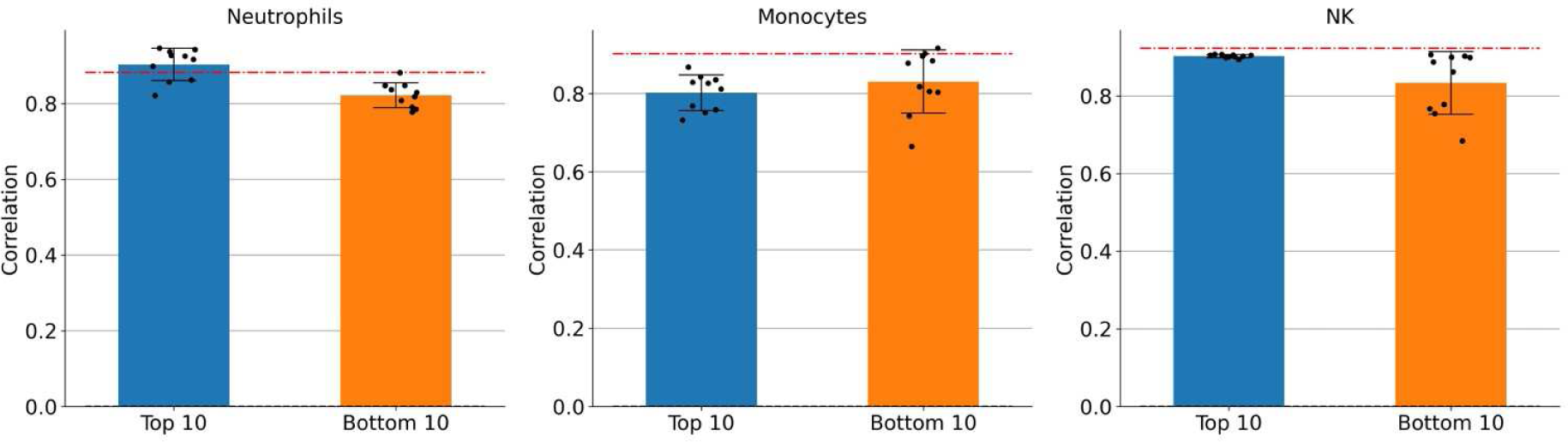
Optimization of reference cell type combinations against the artificial pseudo-bulk dataset. Bar plots showing the difference between the top 10 and bottom 10 Pearson correlations. The red dashed line indicates the baseline when all 13 cell types are considered as reference.

**Figure S17.**
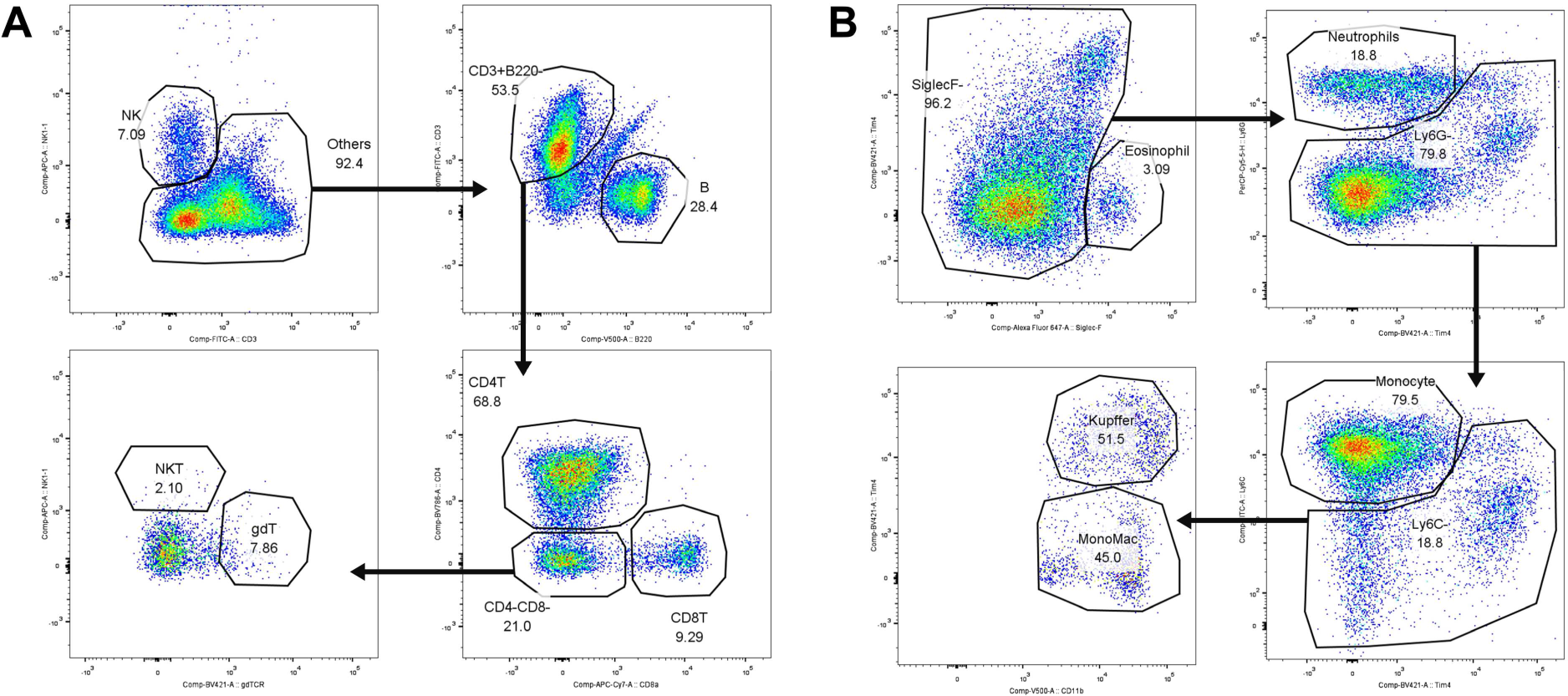
Flow cytometry gating strategy used to reproduce the time-dependent APAP-induced liver injury model described by Bird et al. **A** Lymphocytes: CD4+ T cells (CD4), CD8+ T cells (CD8), gamma-delta T cells (γδT), B cells, natural killer T (NKT) cells, and natural killer (NK) cells. **B** Myeloid cells: monocytes, neutrophils, Kupffer cells, eosinophils, and monocyte-derived macrophages (MDMs).

## Supplementary tables

**Table S1.** Summary of each compound administration.

| **Compound name** | **Route** | **Dose** | **Vehicle** |
| --- | --- | --- | --- |
| alpha-naphthyl isothiocyanate (ANIT) | gavage | 200 mg/kg | 0.5 % methylcellulose |
| acetaminophen (APAP) | gavage | 400 mg/kg | 0.5 % methylcellulose |
| 4,4'-methylene dianiline (MDA) | gavage | 300 mg/kg | 0.5 % methylcellulose |
| thioacetamide (TAA) | gavage | 60 mg/kg | 0.5 % methylcellulose |
| concanavalin A (ConA) | i.v. | 25 mg/kg | 0.9% saline |
| CCl<sub>4</sub> (CCl<sub>4</sub>) | i.p. | 0.5% 10mL/kg | corn oil |
| galactosamine (GAL) | i.p. | 2000 mg/kg | 0.9% saline |

## Supplementary notes

### FUNDING

This work was supported by JSPS KAKENHI Grant-in-Aid for Scientific Research (C) (grant number 21K06663) from the Japan Society for the Promotion of Science, JSPS KAKENHI Grant Number 16H06279 (PAGS), and Takeda Science Foundation.

#### Flow cytometry analysis

##### Liver injury models

Proportions of NK cells (CD45^+^/CD3^−^/NK1.1^+^), NKT cells (CD45^+^/CD3^+^/NK1.1^+^), αβT cells (CD45^+^/CD3^+^/gdTCR^−^), γδT cells (CD45^+^/CD3^+^/gdTCR^+^), eosinophils (CD45^+^/CD11b^+^/Siglec-F^+^), monocytes (CD45^+^/CD11b^+^/Siglec-F^−^/Ly6G^−^/Tim4^−^/Ly6C^+^), monocyte-derived macrophages (CD45^+^/CD11b^+^/Siglec-F^−^/Ly6G^−^/Tim4^−^/Ly6C^−^), neutrophils (CD45^+^/CD11b^+^/Siglec-F^−^/Ly6G^+^/Tim4^−^), and Kupffer cells (CD45^+^/CD11b^+^/Siglec-F^−^/Ly6G^+^/Tim4^+^/Ly6C^−^/F4/80^+^) were acquired via flow cytometry using the following antibodies: PE rat anti-mouse CD45 (ID: 553081), PerCP/Cy5.5 rat anti-mouse CD31 (ID: 562861), FITC rat anti-mouse CD3 (ID: 561798), APC Rat anti-mouse NK1.1 (ID: 550627), BV421 rat anti-mouse γδTCR (ID: 562892), BV605 rat anti-mouse CD11b (ID: 563015), APC-R700 rat anti-mouse F4/80 (ID: 565787), FITC rat anti-mouse Ly6C (ID: 561085), and APC-Cy7 rat anti-mouse Ly6G (ID: 560600). All these antibodies were purchased from BD Biosciences.

##### Time-dependent model of APAP-induced liver injury

Proportions of NK cells (CD45^+^/CD3^−^/NK1.1^+^), B cells (CD45^+^/B220^+^), CD4+ T cells (CD45^+^/CD3^+^/B220^−^/CD4^+^), CD8+ T cells (CD45^+^/CD3^+^/B220^−^/CD4^−^/CD8a^+^), NKT cells (CD45^+^/CD3^+^/B220^−^/CD4^−^/CD8a^−^/NK1.1^+^), γδT cells (CD45^+^/CD3^+^/B220^−^/CD4^−^/CD8a^−^/gdTCR^+^), eosinophils (CD45^+^/CD11b^+^/Siglec-F^+^), neutrophils (CD45^+^/CD11b^+^/Siglec-F^−^/Ly6G^+^/Tim4^−^), monocytes (CD45^+^/CD11b^+^/Siglec-F^-^/Ly6G^−^/Tim4^−^/Ly6C^+^), monocyte-derived macrophages (CD45^+^/CD11b^+^/Siglec-F^−^/Ly6G^−^/Tim4^−^/Ly6C^−^), and Kupffer cells (CD45^+^/CD11b^+^/Siglec-F^−^/Ly6G^−^/Tim4^+^/Ly6C^-^) were acquired via flow cytometry using the following antibodies: PE rat anti-mouse CD45 (ID: 553081), FITC rat anti-mouse CD3 (ID: 561798), BV786 rat anti-mouse CD4 (ID: 563331), APC-C7 rat anti-mouse CD8a (ID: 561967), BV480 rat anti-mouse B220 (ID: 565631), APC rat anti-mouse NK1.1 (ID: 550627), BV421 rat anti-mouse γδTCR (ID: 562892), R718 rat anti-mouse CD45 (ID: 567075), BV480 rat anti-mouse CD11b (ID: 566149), FITC rat anti-mouse Ly6C (ID: 561085), and APC-Cy7 rat anti-mouse Ly6G (ID: 560601), BV421 rat anti-mouse Tim4 (ID: 744874), and PE rat anti-mouse F4/80 (ID: 565410). All these antibodies were purchased from BD Biosciences.

